# Communication subspace dynamics of the canonical olfactory pathway

**DOI:** 10.1101/2024.06.10.598298

**Authors:** Joaquín González, Pablo Torterolo, Kevin A. Bolding, Adriano BL Tort

## Abstract

Understanding how different brain areas communicate is crucial for elucidating the mechanisms underlying cognition. A possible way for neural populations to interact is through a communication subspace, a specific region in the state-space enabling the transmission of behaviorally-relevant spiking patterns. In the olfactory system, it remains unclear if different populations employ such a mechanism. Our study reveals that neuronal ensembles in the main olfactory pathway (olfactory bulb to olfactory cortex) interact through a communication subspace, which is driven by nasal respiration and allows feedforward and feedback transmission to occur segregated along the sniffing cycle. Moreover, our results demonstrate that subspace communication depends causally on the activity of both areas, is hindered during anesthesia, and transmits a low-dimensional representation of odor.

## Introduction

Understanding how different brain areas communicate is a central problem in neuroscience^1,2^. Large-scale cortical connections allow information to be integrated, a process fundamental for sensory perception, decision-making, and cognition^3^. Despite interregional communication being the basis of brain function, major problems challenge its study. At the circuit scale, communication relies on synapses formed by large populations of neurons from different areas, where the number of synapses far exceeds the number of neurons^4^. Moreover, cross-area interactions in the mammalian cortex can be feedforward, feedback, or bidirectional^5–8^ and are often comprised of both excitatory and inhibitory connections.

To understand how brain areas communicate, we need to explain how the neuronal populations in each site interact to allow the transmission of behaviorally-relevant spiking patterns^9,10^. In the neocortex, different areas can interact through a communication subspace^11–18^, a specific region in the state-space (where each dimension corresponds to the activity of a single neuron) of a local population that enables communication. In simpler words, communication occurs through local co-activity patterns that maximize the correlation between two areas. This idea implies that all of the variance and information embedded in the activity of a neural population is not equally transmitted to connected brain regions. Rather, there are specific directions and dimensions in neural activity space along which two regions may co-vary, suggesting that selective information relevant to the function of this specific interregional interaction can be transmitted via this communication subspace. Hence, only the population patterns that align and advance through this subspace would be maximally transmitted between areas.

Olfaction in mammals relies on communication between the olfactory bulb (OB) and the olfactory cortex (PCx; piriform cortex)^19–25^. These areas are densely connected through long-range connections by which different odor features get transmitted. Nonetheless, we still do not know if their populations interact through a communication subspace. Moreover, communication subspaces have been described in the neocortex, raising the question if other cytoarchitectural designs (e.g., paleocortex like the OB and PCx) may be able to support them or which characteristics they may have. Therefore, the study of communication in these regions is likely to offer new insights into the general mechanisms of corticocortical interactions in the brain.

Among the factors influencing communication, temporal coordination of spiking activity plays a major role for interregional interactions. Sniffing patterns impose a natural rhythm to OB and PCx neurons^22,26–32^, coordinating local spiking activity and timing computations from a 300-500 ms window during rest to 100 ms during faster breathing rates^33^. Besides influencing local activity, it is widely recognized that the respiratory rhythm itself influences cognition, improving task performance^34,35^ and regulating cognitive states^36,37^. Consistently, respiration-entrained brain oscillations are prominent in olfactory and associative areas (e.g., hippocampus^38^, prefrontal cortex^39^, somatosensory cortex^40^, amygdala^41^), suggesting that this rhythm may play a role in neuronal communication. Yet, we still do not fully comprehend how this rhythm influences communication at the neuronal firing scale.

Here, we address how population patterns are transmitted in the main olfactory pathway. Our results show that OB and PCx neurons interact through a communication subspace, which supports feedforward and feedback interactions and is entrained by the respiratory cycle, thus revealing how respiration influences spiking transmission. Moreover, experimental manipulations show that subspace activity can be triggered optogenetically and depends on the recurrent PCx connectivity to amplify and filter both OB and PCx responses. From a functional perspective, our results show that this communication channel is abolished during anesthesia and exploited in a flexible way during olfaction to transmit a low-dimensional representation of odor identity.

## Results

### OB and PCx neuronal populations interact through a communication subspace

To understand communication in the main olfactory pathway, we first analyzed simultaneous recordings from OB and PCx populations performed during spontaneous (no odors presented) awake periods in head-restrained mice (Fig. 1A). Because, in principle, any pairwise combination of neurons in each area might form a synapse, we employed canonical correlation analysis (CCA) to study the communication between these populations. Specifically, CCA seeks to maximize the correlation between 2 neuronal populations by finding a communication subspace (CS) in which to project local patterns to yield 2 CS activity signals that are maximally correlated (Fig. 1B; see also Fig S1). In simpler words, the CCA approach identifies which subsets of neurons are correlated between the areas, and the communication subspace of each area is obtained by ascribing large (positive or negative) weights (hereafter referred as CS weights) to these neurons and low weights to uncorrelated neurons; finally, for each region, the CS activity is a signal obtained by the weighted sum of its neuronal activity as dictated by its shared activity axis. Therefore, while the non-weighted summed activity (i.e., multi-unit activity, MUA) of each area may seem unrelated, a subset of correlated neurons (or several subsets) can be identified by the CS framework.

**Figure 1.**
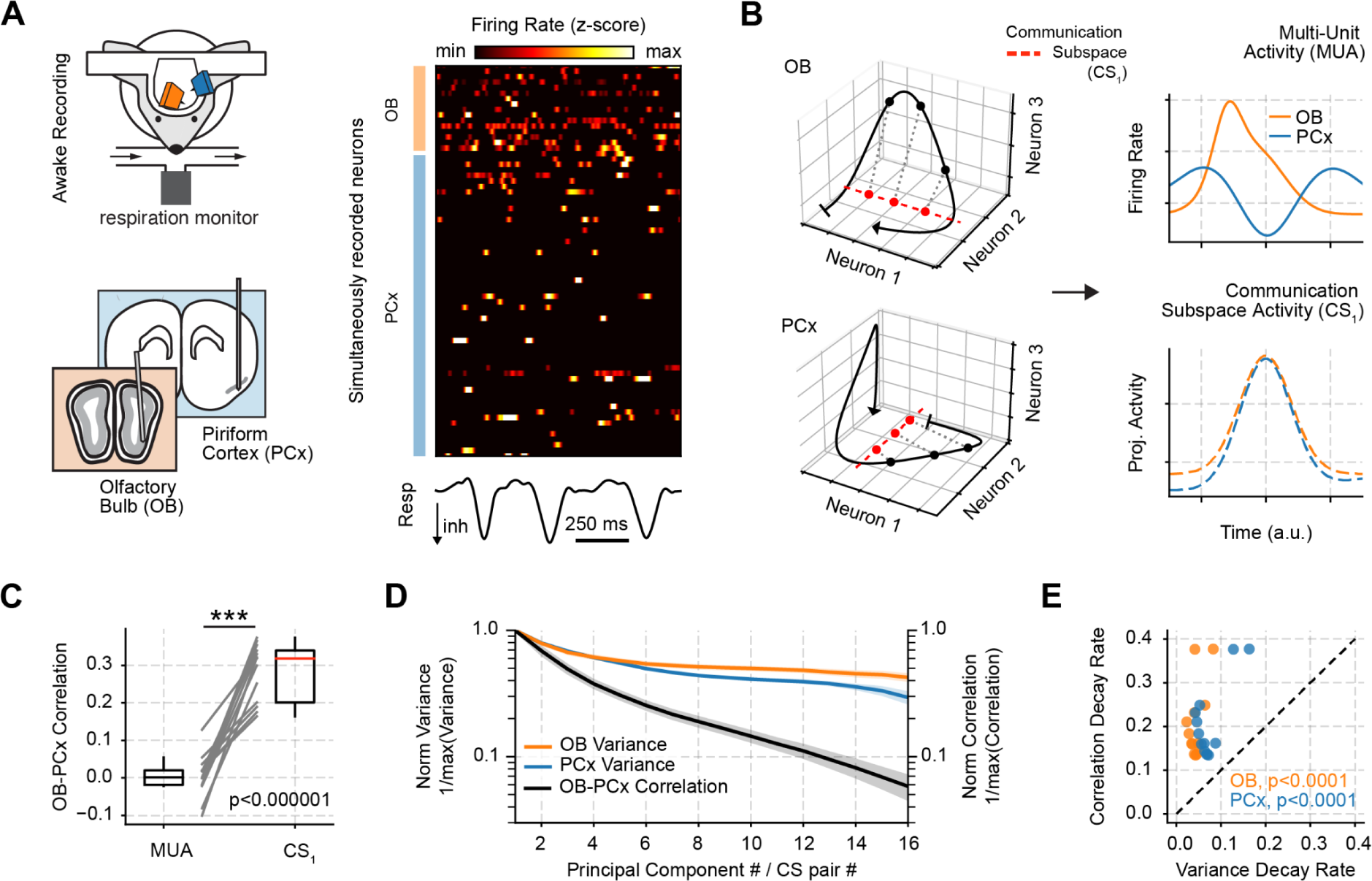
Characterizing the communication subspace between the olfactory bulb and piriform cortex. **A** Experimental recording scheme and probe localization in both olfactory areas (left panels; modified from^22^), and example single unit recordings along with the respiration signal (right). **B** Toy model showing how canonical correlation analysis maximizes cross-area correlations. Canonical correlation achieves this by projecting the local state-space activity (black traces in left panels) to the subspace which maximizes cross-area correlations (CS_1_ red dashed axis). Thus, this method allows to obtain maximally correlated signals across areas (bottom right panel), contrasting with MUA which lacks any correlation (top right panel). **C** OB-PCx Pearson correlation for MUA and CS_1_ signals. Boxplots show the median, 1st, 3rd quartiles, the minimum, and the maximum (excluding outliers); each line shows an individual session. n = 13 recording sessions from 12 mice. **D** Average normalized variance/correlation for each principal component number and CS pair (± SEM; n = 13 recording sessions from 12 mice). For each recording session, variances and correlations were normalized by dividing each value by the maximum variance/correlation. **E** Comparison of the variance and correlation decay for each recording session. The decays were calculated for the normalized variance/correlation as a function of the principal component number/CS pair using a negative exponential fit.

First, we validated the CS framework by studying a toy neuronal model in which we could control communication and local activity. We found that the CS could correctly identify individual subsets of communicating neurons between areas on a background of random local activity (Fig. S1A,B). Moreover, different CS pairs captured the presence of independent linear combinations of neurons (i.e., independent subsets of correlated neurons) by ascribing different sets of weights for each combination (Fig S1C,D). These toy-model results provide a ground-truth reference for the use of CS.

When applying this framework to the OB-PCx data, we first identified how many significant CS pairs exist (Fig. S2). To this end, we compared the correlation coefficient (R) of each CS pair against a surrogate distribution of R values computed by circularly time shifting the spiking activity of one of the populations by random amounts (Fig. S2A,B). For each surrogate, we computed the CS and obtained the R for the first (i.e., most correlated) pair. We repeated this procedure 100 times for each session. We found that the first CS pair (CS_1_) exhibited the largest correlation and that subsequent pairs showed an exponentially decaying distribution of correlation values (Fig. S2C). When comparing against surrogates, only the first four CS pairs showed values above chance for all 13 recording sessions (Fig. S2D). Nevertheless, for the remainder of the paper we focused on the first CS pair since it accounted for the largest correlation.

Notably, our results show that a communication subspace, and not global activity, explains neuronal correlations in the OB-PCx pathway. To contrast CS results, we compared the OB-PCx CS_1_ correlation to the null hypothesis that the communication subspace is equivalent to the population activity (i.e., all neurons considered) in each area. To that end, we computed the MUA signal by summing all single neuron spikes in each area (employing the same time bins as used to compute the CS). Importantly, across all animals and sessions, OB-PCx CS_1_ activity exhibited significantly higher correlations than the OB-PCx MUA (Fig. 1C; t(12) = 3.80, p < 0.000001).

Furthermore, we show that the CS comprises only a “small region” of the state space of each area. To study how local activity compares to the OB-PCx CS, we matched the number of neurons in each area and computed Principal Component Analysis (PCA) and CS. We reasoned that by comparing the decay rates of the variances across principal components to the decay rate of the CS correlations, we can have an idea of how “big” or “small” the communication subspace actually is compared to the local population patterns. To allow comparisons, we normalized the variance of each principal component (by dividing it by the maximum variance across components) and the correlation from each CS pair (by dividing each value by the maximum correlation across pairs). Following this approach, we found that OB-PCx CS correlations decreased much faster than the individual variances of the principal components of each area (Fig. 1D). We quantified the decay rate by a negative exponential fit of the normalized variances and correlations as a function of the component/CS pair number, which confirmed that OB-PCx correlations decrease significantly faster than the local variances (Fig, 1E, OB: t(12) = 3.80, p < 0.000001; PCx: t(12) = 3.80, p < 0.0001).

### Respiration parses feedforward and feedback neuronal interactions

Next, we show that time-lagged interactions characterize CS communication. Since we reasoned that a genuine communication subspace should exhibit a non-zero time lag due to conduction and synaptic transmission delays (Fig. 2A), we computed the CS using firing counts time-lagged between the regions (see also Fig. S3 for a toy model validation), similar to the traditional cross-correlation analysis. Notably, under this analysis, we found that OB leads PCx activity by ∼25 ms, the time lag with the highest correlation between both areas for CS_1_ (Fig. 2B). Thus, hereafter, we obtained all CS_1_ signals by projecting the neuronal activity using the communication subspace calculated using this time delay; this only influences the neuronal weights of each CS_1_ since neither the OB nor the PCx activity is lagged prior to the CCA projection. We further note that CS weights can be computed during a given time window and then applied to project population activity in a different period.

**Figure 2.**
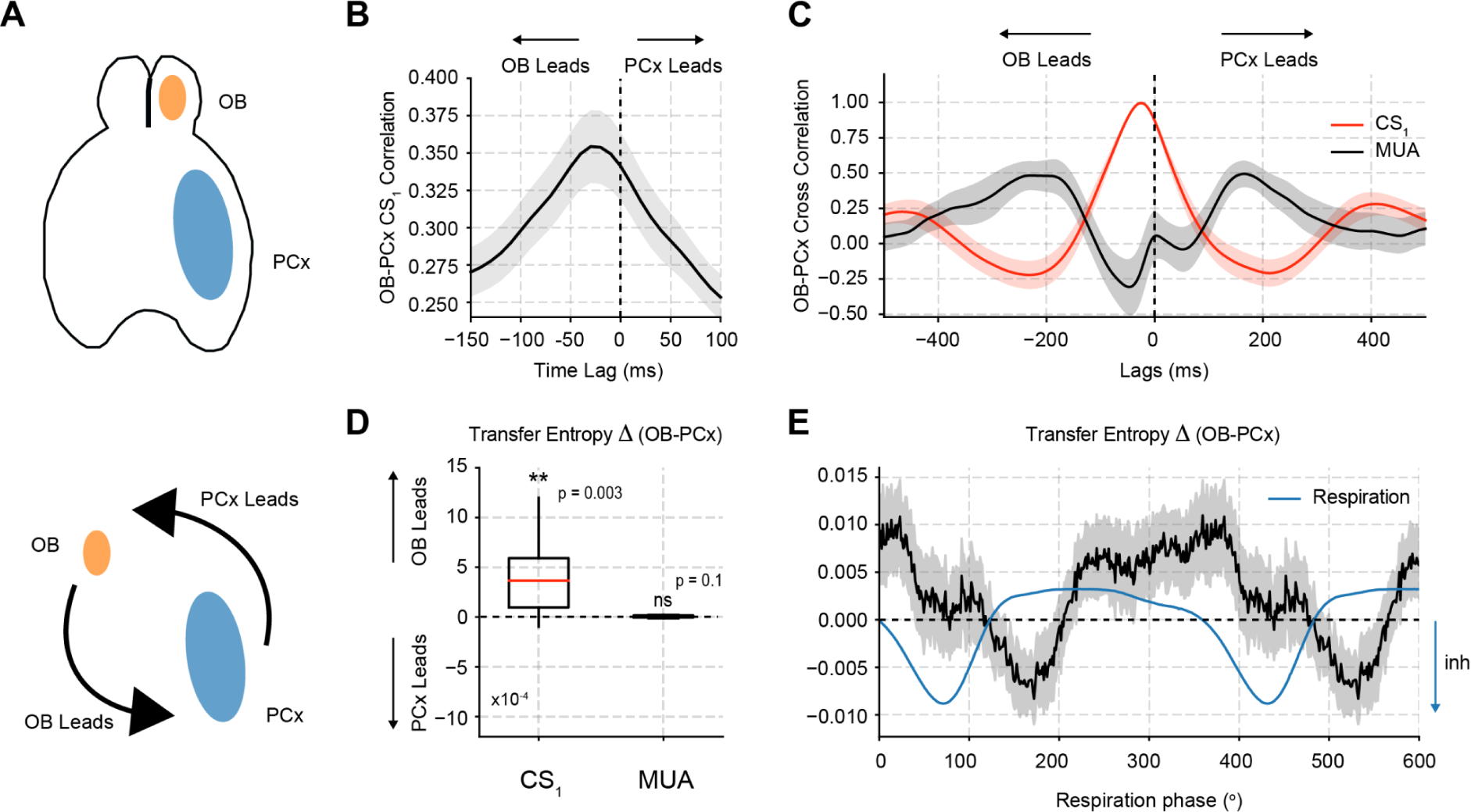
Measuring directionality in the OB-PCx communication subspace. **A** Schematic drawing depicting the OB-PCx pathway and their possible temporal relationships. **B** Average CS_1_ correlation as a function of the time delay between both areas (± SEM; n=13 recording sessions from 12 mice). Notice that by time-lagging one area against the other we find the temporal delay which maximizes the correlation. **C** Average cross-correlogram between both areas for the communication subspace (CS_1_, red) and multiunit (MUA, black) activity (± SEM; n = 13 recording sessions from 12 mice). The cross-correlation for each session was normalized by dividing it by its maximum value. **D** Directionality analysis (Transfer Entropy) between both areas for CS_1_ and MUA. The difference (delta) between OB→PCx and OB←PCx transfer entropy estimates is plotted for each signal. n = 13 recording sessions from 12 mice; ** p < 0.01. **E** Average directionality analysis (Transfer Entropy) as function of the phase of the respiration signal (± SEM; n = 13 recording sessions from 12 mice). As in panel D, the difference (delta) between both directions is plotted. Note a short period of directionality reversal from OB→PCx to OB←PCx at the start of expiration.

To determine whether the communication subspace exhibits directional interactions in the OB-PCx pathway that are not apparent from the global activity, we next computed the OB-PCx CS_1_ cross-correlation (Fig. 2C; red trace) and directionality (Transfer Entropy; Fig. 2D, see Methods) and compared them to the same metrics applied to MUA (black trace). While OB leads PCx activity in the CS_1_ projections, exhibiting a significant OB→PCx directionality (Fig. 2D; t(12) = 3.66, p = 0.003), the MUA results are more challenging to interpret due to the appearance of several peaks in the cross-correlogram and a non-significant directionality (Fig. 2D; t(12) = 1.66, p = 0.12). Of note, the CS_1_ directionality results also hold when computing CS_1_ weights using non-lagged time series (Fig. S4A).

Intrigued by the observation that all CS pairs, on average, exhibited a predominant OB→PCx direction and motivated by the well-described anatomical feedback from PCx→OB, we next investigated whether any evidence of feedback could be observed through CS_1_ activity. To explore this, we divided the respiratory cycle into different phase bins and measured OB-PCx CS directionality in each phase bin. Notably, we observed a reversal in directionality during the course of the respiratory cycle (Fig. 2E). Initially, following the start of inhalation, feedforward interactions predominated (t(12) = 2.86, p=0.01). However, near the start of exhalation, PCx→OB feedback increased and became predominant during a short period (t(12) = −2.65, p = 0.02). Thus, despite the predominance of feedforward directionality, CS_1_ also supports feedback interactions.

### Respiration paces communication subspace activity

We next further characterized respiratory influences and found that, during spontaneous awake periods, the sniffing cycle entrains CS activity. First, we plotted raw and filtered CS_1_ signals along with the nasal respiration recordings and observed each CS_1_ signal to be rhythmically modulated by respiration (Fig. 3A). These results hold across animals (Fig. 3B). Note that the OB to PCx CS_1_ lead was already evident in the raw (solid line), filtered (dashed line; 1-4 Hz), and averaged CS_1_ signals (Fig. 3B top left panel) within each breathing cycle. Consistently, both OB and PCx CS_1_ signals show a prominent 2.5 Hz power spectrum peak (Fig. 3B top right), coinciding with the respiratory peak frequency and coherent to the respiration signal (Fig. 3B bottom). Of note, respiratory entrainment was not restricted to CS_1_ and was also seen for subsequent CS pairs (Fig. S4B). Interestingly, when contrasting CS_1_ and MUA coherence to respiration, we found that both OB and PCx CS_1_ are significantly more coupled to the respiratory activity (Fig. 3B bottom; CS_1_ vs MUA OB-Resp Coherence: t(12) = 6.02, p < 0.0001; PCx-Resp Coherence: t(12) = 5.12, p = 0.0002), suggesting that respiration preferentially entrains subspace activity.

**Figure 3.**
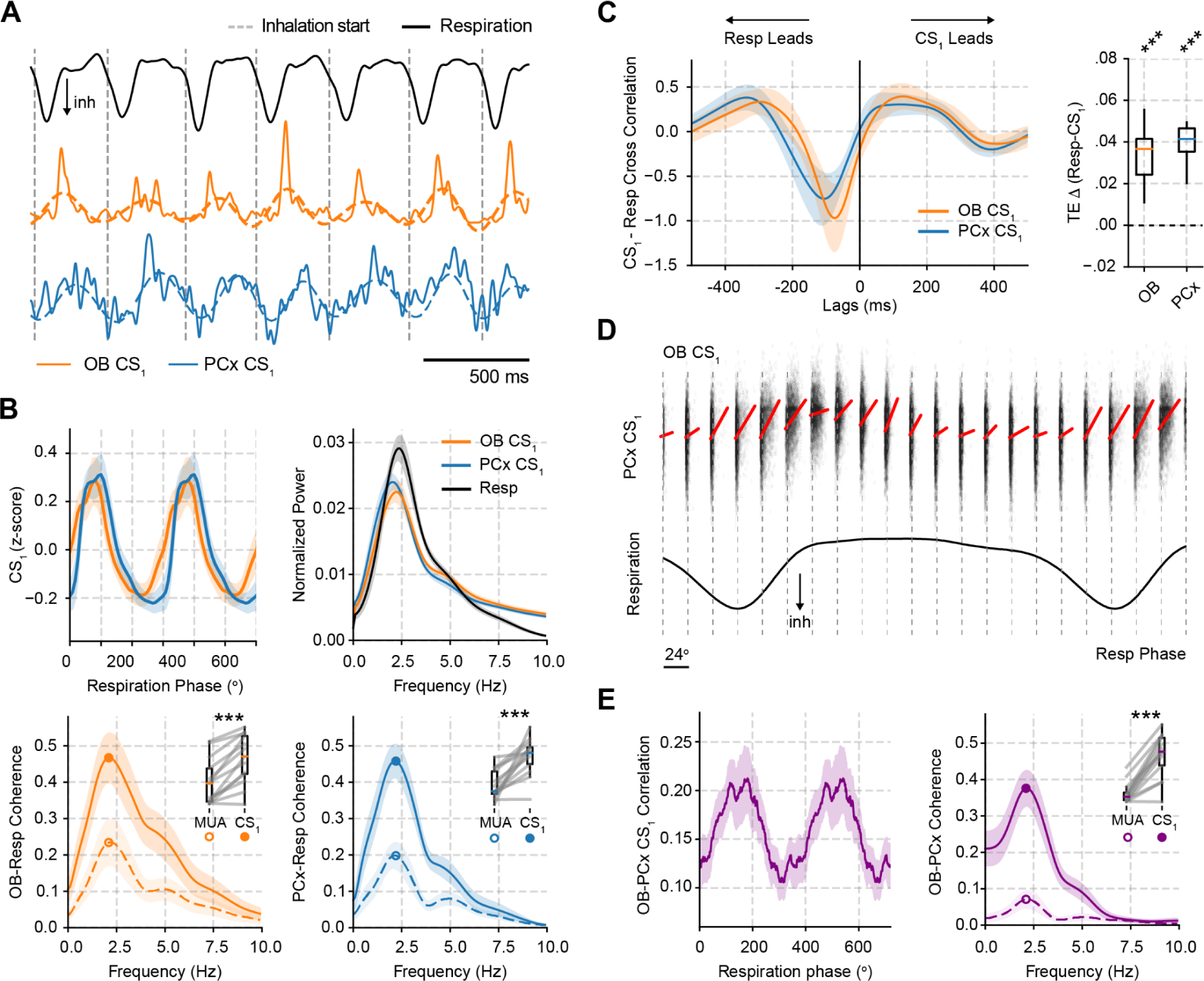
Respiration drives the activity of the communication subspace. **A** Left: Example of OB and PCx CS_1_ along with the respiration signal. Dashed vertical lines indicate inhalation starts. Dashed colored lines show the 1-4 Hz filtered CS_1_ for each area. **B** Top Left: Average CS_1_ as a function of respiratory phase (± SEM; n = 13 recording sessions from 12 mice). Top Right: Average power spectrum of the CS_1_ and respiration signals (± SEM). Bottom: Average CS_1_ and MUA coherence to respiration (± SEM). Insets: Boxplots comparing peak CS_1_ and MUA coherence to respiration (frequency indicated by the dot); each line shows an individual mouse. *** p < 0.001. **C** Left: Average cross-correlogram between respiration and OB/PCx CS_1_ (± SEM; n = 13 recording sessions from 12 mice). Right: Directionality analysis (Transfer Entropy) between respiration and OB/PCx CS_1_. The difference (Δ) between Resp→CS_1_ and CS_1_→Resp transfer entropy estimates is plotted for each signal. **D** OB-PCx CS_1_ correlation as a function of the respiratory phase. The top panel shows an example correlation from a single recording session; the red lines show best linear fit for each respiratory phase. Below we show for the same recording session the average respiratory waveform. **E** Left: average CS_1_ correlation (± SEM; n = 13 recording sessions from 12 mice) during the respiratory cycle. Right: Average CS_1_ and MUA OB-PCx coherence (± SEM). Inset: Boxplots comparing CS_1_ and MUA in the frequency indicated by the dot. *** p < 0.001.

We next show that respiration drives CS_1_ activity. We started by computing cross-correlograms between OB/PCx CS_1_ signals and respiration. Importantly, we found that the cross-correlation function was not symmetrical and was heavily skewed towards negative lags, corresponding to respiration leading CS_1_ activity in both areas. Note that, consistent with the previous OB-PCx directionality results, the OB CS_1_ exhibited a shorter lag in the respiration cross-correlogram compared to the PCx CS_1_.These results were confirmed through transfer entropy analysis showing significant Resp→CS_1_ results for both OB and PCx (Fig. 3C, right; OB: t(12) = 6.19., p < 0.0001; PCx: t(12) = 10.39, p < 0.000001).

Besides finding respiratory entrainment of local CS signals, we further investigated whether OB-PCx correlations depend on the phase of the breathing cycle. This idea implies that maximal communication occurs during a specific phase of the respiratory cycle. To test this hypothesis, we binned the spontaneous CS_1_ activity according to the respiratory phase and measured the OB-PCx CS_1_ correlation in each phase bin (using different respiratory cycles as samples; Fig. 3D). We found a significant modulation of the CS_1_ correlations (Z(12) = 4.26, p = 0.01, Rayleigh test); in particular, correlations increased following the start of inspiration and then declined in the later part of the respiratory cycle (Fig. 3E). Consistent with this result, OB-PCx CS_1_ coherence showed a synchronization peak at the respiratory frequency (Fig. 3E). Moreover, OB-PCx CS_1_ coherence was significantly higher than OB-PCx MUA coherence (Fig. 3E inset; OB-PCx Coherence: t(12) = 7.97, p < 0.0001), further confirming that OB and PCx are likely to interact through a communication subspace and not through the global population.

### CS membership relates to spiking phase preference during the respiratory cycle

Next, we show that CS_1_ weights relate to the spiking phase preference of single neurons during the respiratory cycle. We plotted the average activity of all putative principal cells in each area as a function of the respiratory phase and sorted them according to their CS_1_ weights (Fig. 4A). We found that most OB neurons exhibited a single preferred respiratory phase near the inhalation start (40-140°), and that the most positive PCx neurons (+CS_1_ weights) showed a similar phase preference to OB neurons. Nevertheless, the remaining PCx neurons (-CS_1_ cells) exhibited an anti-phase pattern (that is, preferred the opposite phase than OB neurons or PCx +CS_1_ cells). Moreover, the z-scored firing during the 40-140° respiratory phase significantly correlated with the CS_1_ weights in both areas (OB: R = 0.23, p < 0.0001; PCx: R = 0.34, p < 1×10^−20^). Note that while +CS_1_ cells in either region exhibited similar firing levels during this phase bin, −CS_1_ cells fired close to baseline in the OB and decreased their average spiking in the PCx (Fig. 4A). Consistently, the distribution of CS_1_ weights was skewed toward negative values in the PCx, corresponding to the large majority of (likely inhibited) anti-phase neurons, and was significantly different from the CS_1_ weight distribution of OB (D(110) = 0.12, p < 0.0001, Kolmogorov-Smirnov test). Importantly, we found a significant correlation between the absolute CS_1_ weights and the respiratory coupling strength of single cells (Fig. 4B, OB: R = 0.41, p < 1×10^−13^; PCx: R = 0.37, p < 1×10^−29^), suggesting that respiration preferentially entrains communicating neurons in both areas.

**Figure 4.**
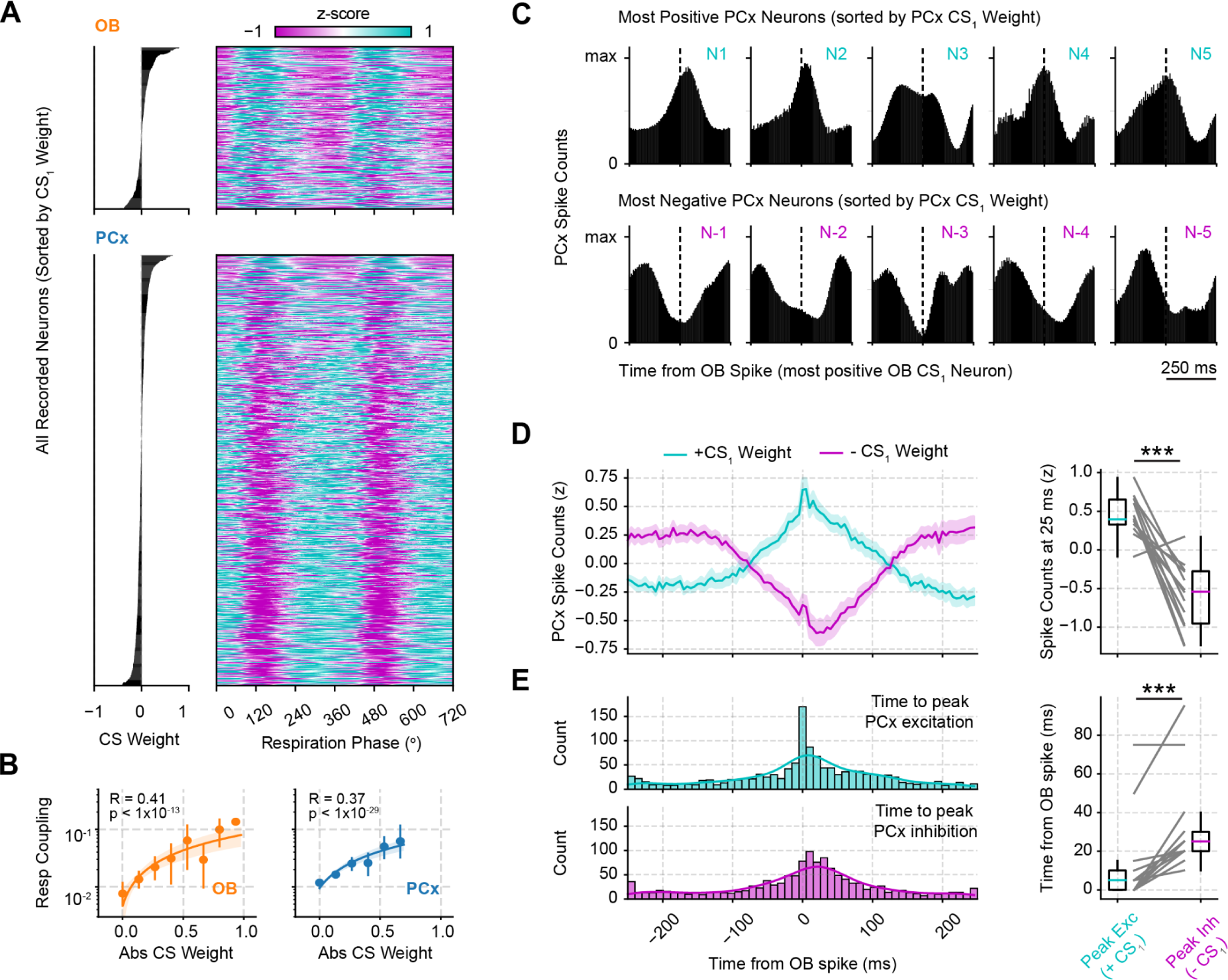
Excitatory-inhibitory spike relations characterize OB-PCx CS_1_ communication. **A** Neuron spiking during each phase bin of the respiration cycle (right; 0 degree corresponds to the start of the inhalation); neurons are sorted according to the CS_1_ weights (shown on the left panel). OB: 320 cells; PCx: 849 cells. **B** Correlation between absolute CS_1_ weights and the respiratory coupling strength (Modulation Index) of single cells. Note that bins were only employed for representational purposes. Error bars show 95% CI. **C** Spike-spike cross-correlations between a single OB neuron (largest CS_1_ weight) and 10 PCx neurons (top 5 most positive [top row] and negative [bottom row] CS_1_ weights). **D** Average spike-spike cross-correlogram (± SEM; n = 13 recording sessions from 12 mice) for the most positive OB vs. most positive (teal) and most negative (magenta) PCx CS_1_ weighted neurons. Boxplot: Activity at the +25 ms time delay; each line shows an individual mouse. **E** Histogram showing the distribution of maximum excitation (teal) and maximum inhibition (magenta) times for the 10 most positive and 10 most negatively CS_1_ weighted neurons across animals. Boxplot: Average time to maximum excitation/inhibition of positive/negative CS_1_ weighted PCx neurons across animals. *** p < 0.001.

We next show that putative excitatory and inhibitory interactions explain CS_1_ weights. First, we selected the most positive CS_1_ OB neuron from a representative recording and analyzed the spike counts of the top five +CS_1_ and −CS_1_ PCx cells. We found that the +CS_1_ PCx cells tended to spike shortly after OB spikes, while −CS_1_ PCx cells tended to be inhibited (Fig. 4C). This result was consistent across animals when analyzing the top ten +CS_1_ OB neurons against the top ten +CS_1_ and −CS_1_ PCx neurons (Fig. 4D, t(12)=7.38, p < 0.001). Note that inhibition occurred later compared to excitation in the PCx (t(12) = 2.25, p < 0.001; Fig. 4E), suggesting that inhibition, on average, may take more synaptic steps than excitation. These results suggest that subspace interactions and respiratory firing phase preferences are likely to be intrinsically related.

### OB-PCx communication subspace interactions depend on intact synaptic connectivity

We next demonstrate that direct communication underlies OB-PCx subspace interactions. To this end, we analyzed CS_1_ activity during optogenetic stimulation of OB excitatory cells (Fig. 5A). For these analyses, we calculated CS_1_ weights employing the laser stimulation periods. We found that light stimulation caused a large amplitude, transient CS_1_ response (Fig. 5B). The transient response first occurred in the OB CS_1_ activity and appeared after a short <10-ms latency in the receiving PCx population (Fig. 5B; Time to CS_1_ peak: t(4) = 5.65, p = 0.004), confirming the predominant OB-PCx communication directionality. In the OB, the transient response was followed by a lower amplitude sustained activity lasting through the entire stimulation period (Fig. 5B). Noteworthy, the spontaneous activity of the cells identified during laser stimulation also exhibited the phase-antiphase CS_1_ spike pattern segregation within respiratory cycles (Fig. S5) as found in our earlier analysis (c.f. Fig. 4A).

**Figure 5.**
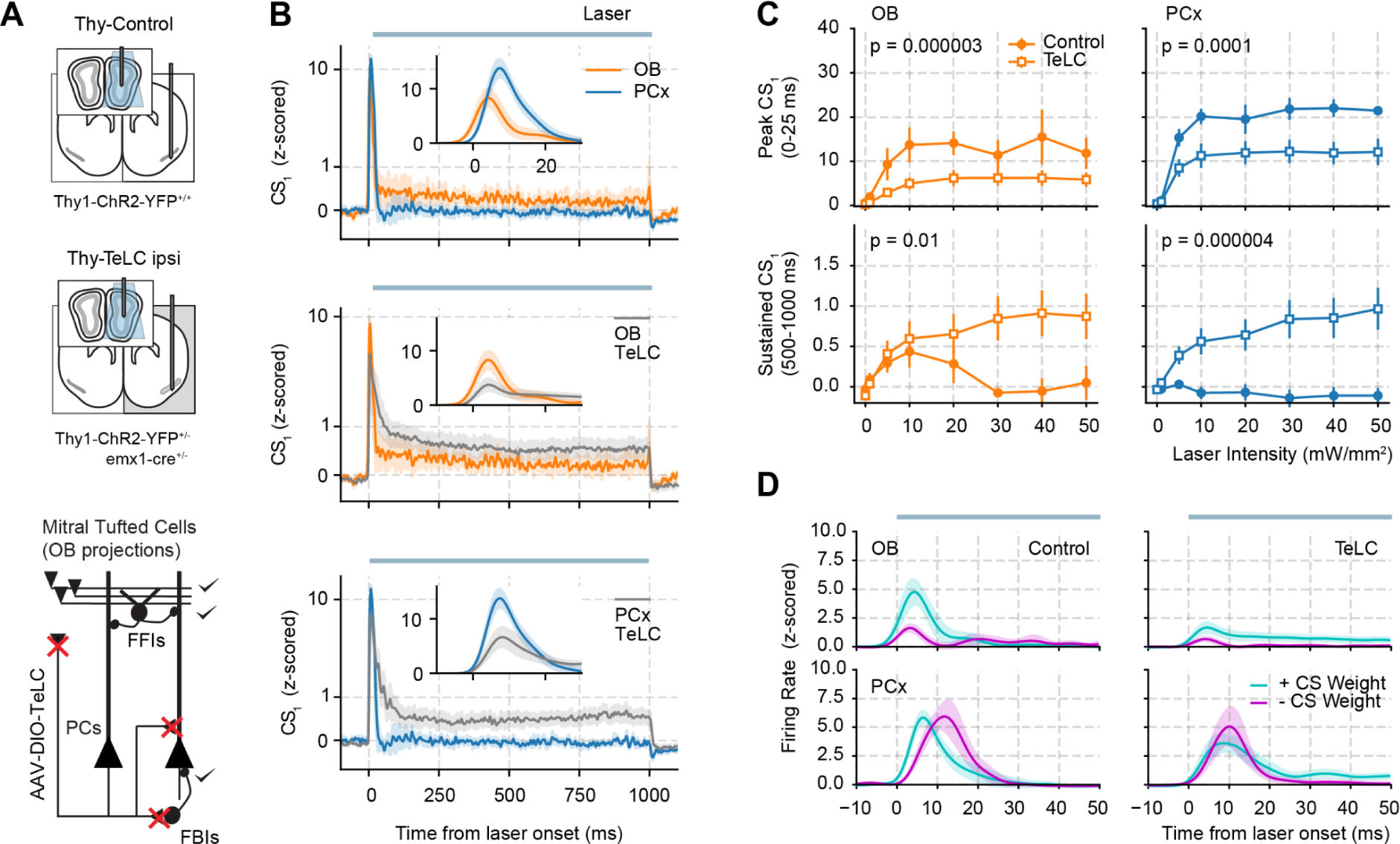
Probing the mechanisms behind the OB-PCx communication subspace. **A** Top: Experimental recording conditions for each group. OB and PCx recordings were made from the hemisphere containing TeLC expression (modified from^22^) during optogenetic stimulation of the OB (laser duration = 1 s). Bottom: Schematic of circuit changes after TeLC expression in principal cells (PCs) of the piriform circuit (MTCs: mitral cells; FFIs: feedforward interneurons; FBIs: feedback interneurons). TeLC expression blocks synapses from PCx principal cells (red crosses). **B** Average OB and PCx CS_1_ activity triggered by optogenetic stimulation of OB projecting neurons (± SEM; Control: n = 5 recording sessions from 5 mice; TeLC: n = 14 recording sessions from 7 animals). Insets highlight the initial response. **C** Average CS_1_ activity (± SEM) as a function of the laser intensity). The laser stimulation period was divided into an initial (0-25 ms) and a sustained response (500-1000 ms). **D** Triggered spiking activity of “positive” and “negative” cells according to their CS_1_ weights. The most positive and most negative cell from each session was employed here. The panel shows a zoom-in view of the first 50 ms following laser onset.

Next, we show that 1) recurrent excitation within the PCx amplifies the initial (transient) CS_1_ response, 2) lateral inhibition shuts down the sustained component of the CS_1_ response, and 3) feedback projections from the PCx influence OB CS_1_ activity. To probe these mechanisms, we compared control animals with animals expressing TeLC in the PCx. TeLC expression blocks all synapses from PCx principal cells while preserving principal cell excitability (i.e., from afferent inputs)^22^, allowing us to probe how local PCx interactions shape the CS_1_ activity in response to optogenetic stimulation of the OB (Fig. 5A). Notably, we found that the initial, transient CS_1_ response (0-25 ms) decreased in both areas of TeLC-infected animals (Fig. 5B,C). Interestingly, however, the sustained CS_1_ response (500-1000 ms) increased significantly also for both areas in the TeLC group (Fig. 5C; note that the sustained response is nonexistent in the PCx of control animals). Hence, these results causally demonstrate that OB-PCx subspace communication involves both feedforward (OB→PCx) and feedback (PCx→OB) projections, with the latter conclusion inferred from the fact that altering PCx connectivity alters OB CS_1_ activity.

We next compared the activity of the most positive and negative CS weighted cells following OB stimulation (Fig. 5D). In control animals, +CS_1_ OB cells fired more than −CS_1_ cells during the first 25 ms (t(4) = 4.08, p = 0.01). In addition, +CS_1_ PCx cells fired earlier than −CS_1_ cells (t(4) = 3.98, p = 0.01). In TeLC recordings, +CS_1_ and −CS_1_ OB cells showed a similar firing rate difference as in control animals (t(13) = 2.14, p = 0.04). On the other hand, the time difference previously observed for +CS_1_ and −CS_1_ PCx cells disappeared in TeLC animals (t(13) = 0.17, p = 0.86). Thus, these results suggest that +CS_1_ PCx cells actively recruit inhibition and suppress −CS_1_ spiking during the first 10 ms of OB laser stimulation, leading to their different spike time-courses, and that, consequently, this effect is lost in TeLC recordings due to the lack of feedback inhibition recruitment.

### Anesthesia disrupts OB-PCx subspace communication

Next, we show that subspace communication depends on the cognitive state and, consequently, is altered during anesthesia. First, we compared CS_1_ correlations between awake and ketamine/xylazine anesthesia recordings. For this analysis, we computed CS_1_ weights during awake periods and used them to project neuronal activity during anesthesia (Fig. 6A). We found that CS_1_ correlations significantly decreased during loss of consciousness (Fig. 6B left; t(9) = 4.44, p = 0.001), confirming that the subspace that allowed for communication during wakefulness no longer does during anesthesia. Thus, this result suggests a cognitive role for CS_1_ communication.

**Figure 6.**
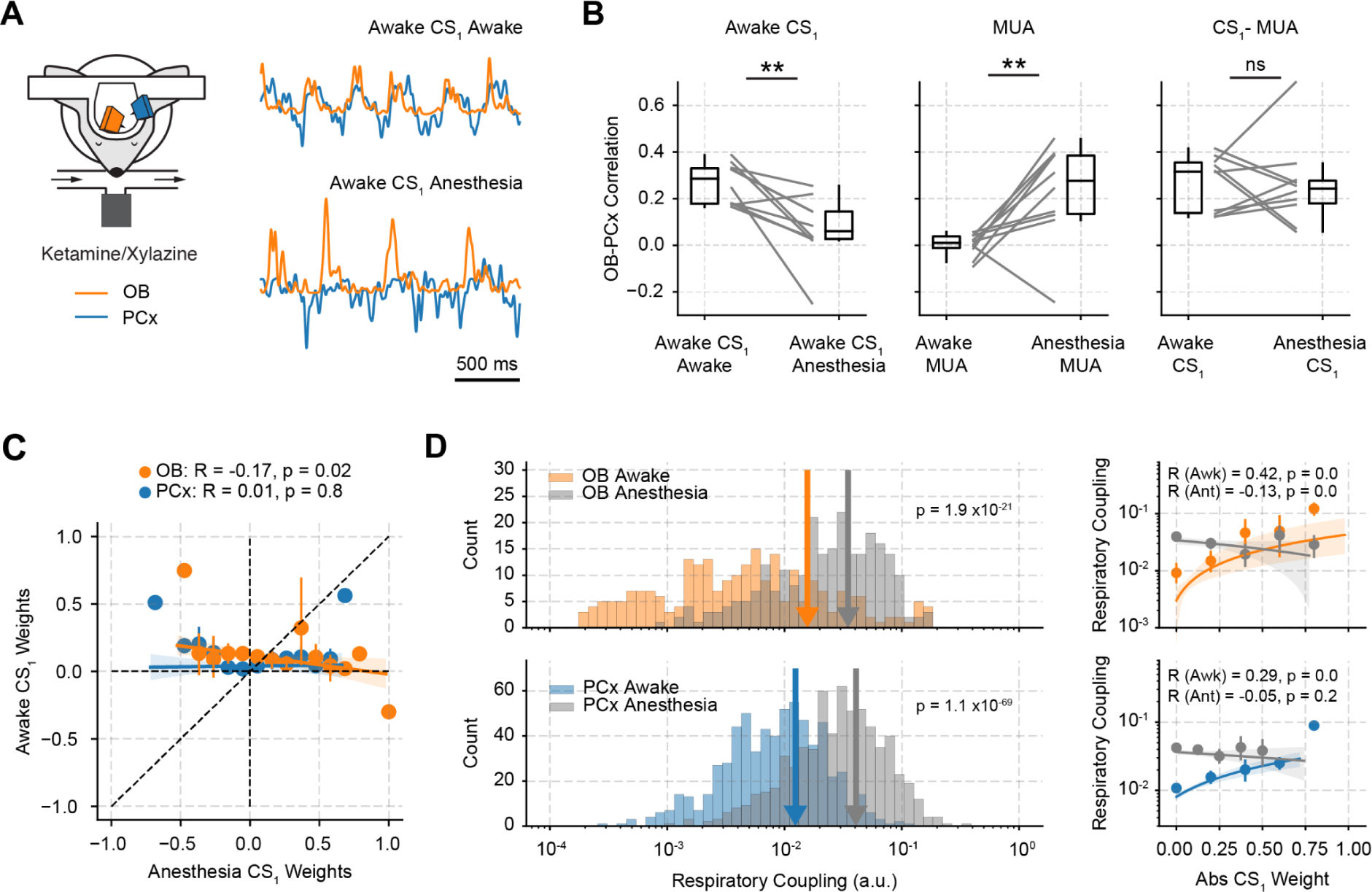
Ketamine-xylazine anesthesia alters CS_1_ communication. **A** Example CS_1_ traces during head-fixed wakefulness and after ketamine-xylazine anesthesia. CS_1_ weights were computed during awake periods and later used to project neuronal activity during anesthesia. **B** Left: OB-PCx CS_1_ correlation during awake and anesthesia periods. CS_1_ weights were computed during awake periods and later used during anesthesia, as in the previous panel. Right: OB-PCx CS_1_ correlation, but with anesthesia CS_1_ weights computed during anesthesia. n = 10 recording sessions from 9 mice. Of note, since OB-PCx correlations largely increase during anesthesia, CS_1_ correlations were normalized by subtracting the OB-PCx MUA correlation. **C** Correlation between CS_1_ weights computed exclusively during awake and anesthesia periods. **D** Left: Distribution of respiratory coupling strength (Modulation Index) for all single cells during awake and anesthesia recordings. The vertical arrows show the distribution mean for each case. Right: Correlation between absolute CS_1_ weights and the respiratory coupling of single cells. Note in C and D that bins were only employed for representational purposes. Error bars show 95% CI.

Moreover, we analyzed how neuronal communication occurs during anesthesia and found that a new CS emerges unrelated to the awake channel. For these analyses, we computed CS weights exclusively using neuronal activity during anesthesia and compared them to the awake subspace weights. Interestingly, we found that both subspaces accounted for similar correlation gains above MUA correlations (Fig. 6B right; t(9) = 0.13, p = 0.89). This control is important since MUA correlations greatly increased during anesthesia (Fig. 6B middle; t(9) = 3.76, p = 0.004). Notably, anesthesia and awake CS_1_ weights were not positively correlated (Fig. 6C, OB: R = −0.17, p = 0.02; PCx: R = 0.01, p = 0.8), confirming that different groups of neurons communicate within each state. Moreover, despite respiratory entrainment of single neurons significantly increasing during anesthesia (Fig. 6D, left panel, OB: t(247) = 10.46, p = 1.9 ×10^−21^; PCx: t(652) = 20.0, p = 1.1 ×10^−69^), the CS_1_ weights were not positively correlated with the level of respiratory coupling in this state (Fig. 6D, right panel, OB: R = −0.13, p = 0.04; PCx: R = −0.05, p = 0.22), contrasting therefore with the results obtained for the awake CS_1_ (c.f., Fig. 4B). The lack of positive correlation between CS weights and spike-respiration coupling during anesthesia (despite the overall coupling increase) demonstrates that respiratory CS entrainment is not a mere consequence of high respiratory modulation of single cells, and further suggests a cognitive role for the differential respiratory entrainment of communicating cells.

### Olfactory communication occurs through a general and a specialized channel

We investigated the functional role of the OB-PCx CS and show that communication during olfaction occurs through a general (“spontaneous” CS_1_) and a specialized channel (“odor” CS_1_). We took two complementary approaches in these analyses: we derived CS weights exclusively during either (1) spontaneous periods or (2) during odor sampling. We found that spontaneous CS_1_ and odor CS_1_ weights were weakly but significantly correlated (OB: R = 0.3, p < 1×10^−7^; PCx: R = 0.3, p < 1×10^−18^; Fig. S6A,B). Prior to analyzing group data of CS_1_ activity, we multiplied all CS_1_ traces by the sign of their response to odors; this was necessary since two animals exhibited CS_1_ activity decreases to odors in both areas (Fig.S7) and we wanted to study changes in the magnitude of CS_1_ activity. When analyzing odor and odorless sniffs, we found an increase in both the spontaneous and odor CS_1_ activity magnitude during odor sampling in both areas (time-stamps marked by blue and orange dots in Fig. 7B; p<0.05, one-sided t-test). Note that odor CS_1_ showed a more sustained odor-evoked response (0-200 ms) compared to the spontaneous CS_1_ (Fig. 7B).

**Figure 7.**
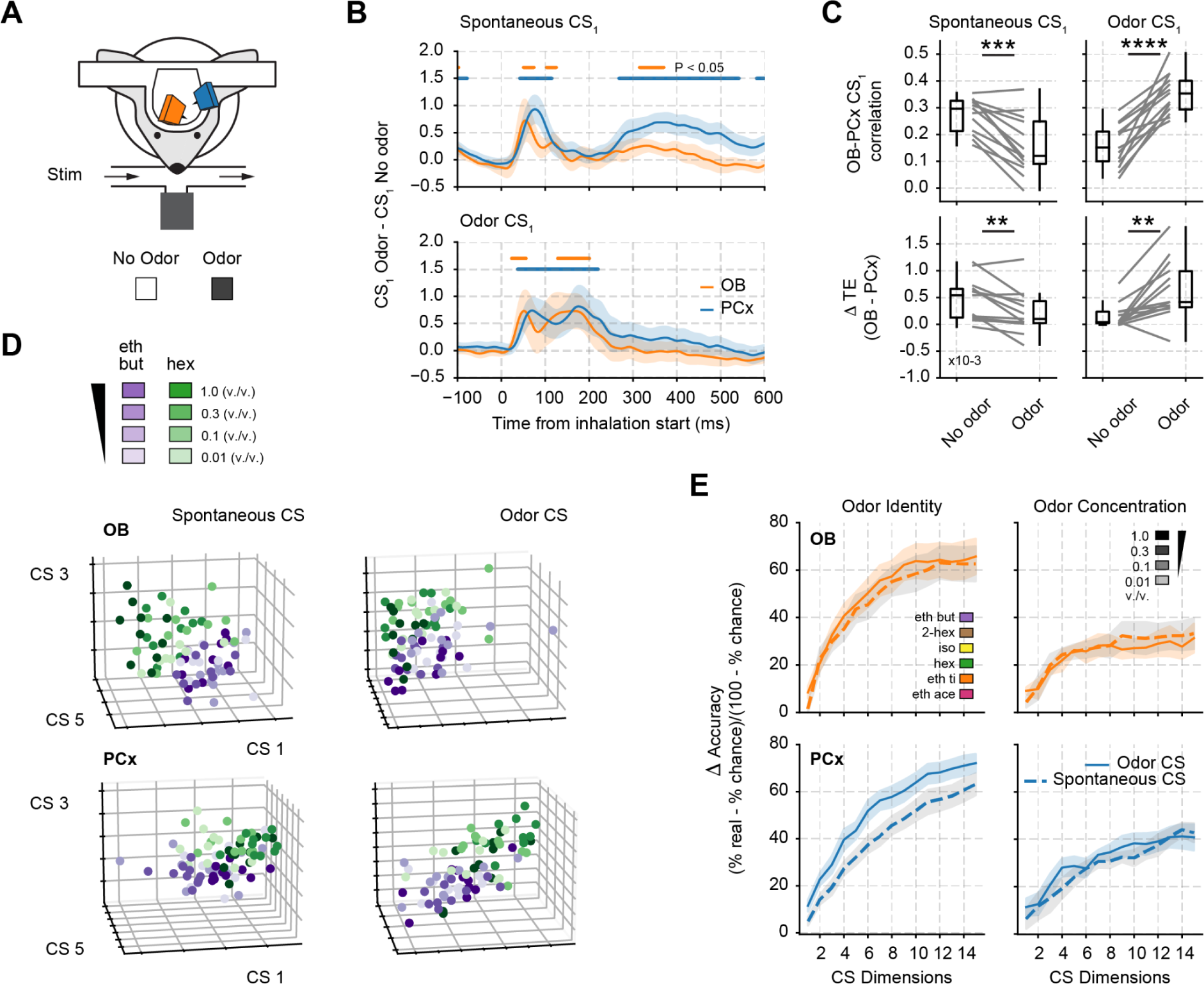
The OB-PCx communication subspace allows the transmission of odor identity. **A** Experimental recording scheme and odor panel employed in the experiments (modified from ^22^). **B** Average CS_1_ activity difference (between odor and odorless sniffs) triggered by inhalation start (± SEM; n = 13 recording sessions from 12 mice). **C** Top: OB-PCx CS_1_ correlations after odor and no odor sniffs (n = 13 recording sessions from 12 mice). Bottom: Directionality analysis (Transfer Entropy) between both areas for CS_1_ during odor and odorless sniffs. The difference (delta) between OB→PCx and PCx→OB transfer entropy estimates is plotted for each condition. **D** Example CS activity for 3 CS pairs from a representative animal obtained during the presentation of 2 different odorants at different concentrations. Each dot shows the average CS activity (0-500 ms following inhalation start) during an individual trial. Note the clustering of purple shaded vs green shaded points, indicating odor separation. **E** Average identity/concentration decoding accuracy (± SEM; n = 13 recording sessions from 12 mice) as a function of the number of aggregated CS pairs (“CS dimensions”) considered for the analysis. The data employed consisted of the average CS activity (for each dimension) during the first 500 ms following inhalation start. For all panels, CS_1_ weights were computed either during exclusive no odor sniffs (spontaneous CS_1_) or exclusively during odor sniffs (odor CS_1_).

Notably, we observed a functional switch in CS communication during odor presentations. Namely, OB-PCx spontaneous CS_1_ correlations decreased during odor presentations compared to odorless periods (Fig. 7C and S6; t(12) = 5.29, p = 0.00019), while odor CS_1_ correlations increased (Fig. 7C, t(12) = 7.21, p = 0.00001). These changes were accompanied by a significant decrease in OB→PCx directionality during odor sampling for the spontaneous CS_1_ (Fig. 7C; t(12) = 3.29, p = 0.006) and a significant increase in OB→PCx directionality for the odor CS_1_ (Fig. 7C; t(12) = 3.84, p = 0.002).

We next analyzed spontaneous CS_1_ activity in response to increasing concentrations of a single odorant (Fig. S8A). Our results showed that for both spontaneous and odor CS_1_, OB activity increased with odorant concentration, while PCx did not (Fig. S8A). Nonetheless, spontaneous OB-PCx CS_1_ correlations decreased as concentration increased, while odor CS_1_ correlations did not depend on concentration (Fig. S8B). Thus, these results further suggest that communication during olfaction involves a mixture of a common (concentration-dependent) and a specialized (concentration-independent) channel.

### Odor identity is transmitted through the communication subspace

Finally, we demonstrate that the OB-PCx CS transmits odor identity as a low-dimensional representation. We analyzed the activity of several CS pairs since odors are likely represented by different neuronal combinations spanning multiple CS pairs (recall that independent linear combinations give rise to different CS pairs; Fig. S1C,D). When plotting the average amplitude of three CS pairs for the presentation of 2 different odorants (0-500 ms after inhalation, 0.01, 0.01, 0.3, 1.0 v./v. concentrations), we found that regardless of concentration, each odor formed a distinct cluster in this low-dimensional subspace (Fig. 7D). Similar results held for 6 odorants at equal concentrations (Fig. S9). The intersection of clusters suggests that the transmission of odorant identity occurs through overlapping, not independent, combinations of neurons (Fig. S10; see also Supplementary Discussion).

Subsequently, we trained and tested a linear decoder employing all available odor presentations to either decode identity from a six-odorant panel at 0.3 v./v. concentration or decode concentration from a two-odorant panel at 0.01, 0.1, 0.3, 1.0 v./v.. As shown in Figure 7E, we successfully decoded identity and concentration above chance levels (1/6 for odors, 1/4 for concentrations) for all the increasing combinations of CS pairs (i.e., CS_1_, [CS_1_,CS_2_], [CS_1_,CS_2_,CS_3_], etc; p<0.01 for all dimensions). Comparing identity and concentration decoding (note the normalization of accuracies to their chance levels), we found that CS activity was significantly more specific to odor identity than concentration (OB Odor CS: F(1,14) = 93.2, p < 10^−18^; OB Spontaneous CS: F(1,14) = 64.13, p < 10^−13^; PCx Odor CS: F(1,14) = 133.1, p < 10^−24^; PCx Spontaneous CS: F(1,14) = 49.1, p < 10^−10^).

Importantly, when comparing spontaneous and odor CS decoding accuracies, we found for the PCx that its odor CS provided significantly better decoding results than the spontaneous CS (p<0.05 for all dimensions), confirming that the olfactory cortex likely employs both a general and a specialized channel for representing odor.

## Discussion

Here, we addressed how population patterns are transmitted in the main olfactory pathway. Our results unveil that a low-dimensional shared subspace (i.e., a communication subspace) forms the major communication channel between OB and PCx neuronal populations. Notably, this communication subspace is paced by nasal respiration and allows feedforward and feedback interactions to occur in different phases of the sniffing cycle. We show that subspace activity can be commanded by OB optogenetic stimulation and that it depends on lateral excitation and inhibition in the PCx to shape initial and sustained responses.

From a functional perspective, our results suggest a cognitive role for CS communication since 1) anesthesia disrupts CS communication, 2) subspace activity increases during odor sampling, and 3) this communication channel transmits a low-dimensional representation of odor identity. This low-dimensional representation carries more identity than intensity information, consistent with previous experimental results,^22^ though concentration information could still be decoded above chance from CS activity, also consistent with experimental and modeling results.^23,42^

### CS analysis offers a comprehensive view of neuronal interactions in the OB-PCx pathway

Several factors complicate our understanding of the olfactory system and how its neurons communicate and represent smell across all major olfactory hubs. One significant challenge is its spatial heterogeneity. Olfactory sensory neurons (OSNs) located in the olfactory epithelium, each expressing a single olfactory receptor, converge into functional units known as glomeruli inside the OB^43^. Despite the order in OSN projections, OB glomeruli lack a detailed chemotopic structure and only exhibit some coarse organization.^44^ Mitral and tufted cells (MTCs) contact these glomeruli and project to other olfactory centers, sending spatially diverse projections, principally to the PCx.^19,20,22,25,45–49^ Consequently, the PCx inherits this lack of a chemotopic map, meaning that the same odorant can activate neurons in distant sites of the same area.^20,28,45,46,50–52^

To identify coordinated neuronal assemblies across these areas, we analyzed the OB-PCx CS, similar to recent efforts in the visual and motor systems.^11–14^ This analysis allowed us to pinpoint groups of cells across areas that exhibit maximal correlation in time and form functional partnerships following each sniff. Crucially, by examining the CS weights of each neuron, we could differentiate between putative excitatory and inhibitory relationships occurring during the respiratory cycle. Although CS weights could be attributed to respiratory phase preferences, the inactivation of the local piriform recurrent circuitry demonstrated that local inhibition causes the firing pattern differences between +CS and −CS cells. Therefore, our results suggest that subspace interactions and the respiratory firing phase preference of single cells are likely to be intrinsically related.

### Respiration drives neuronal communication

It has become increasingly clear that the respiratory rhythm influences the activity of a wide variety of brain areas – well beyond olfactory centers.^27,33,34,38,40,41,53–57^ This rhythm, impacting faster local oscillations,^26,27^ is proposed to play a role in integrating information across the brain^33,58^. Our findings provide direct support for this notion by demonstrating at the neuronal level that: 1) respiration exerts a more pronounced modulation on local communication subspace (CS) activity compared to multi-unit activity (MUA), 2) absolute CS weights predict respiratory entrainment of individual cells during awake recordings, and 3) respiration modulates synchronization between the OB and PCx within the communicating populations. These results highlight the direct impact of this body rhythm on neuronal communication in the main olfactory pathway, suggesting it may also serve similar functions in other upstream areas.

How does respiration influence interregional communication? Our study suggests that it does so by modulating specific directions in neural state-space that define the CS. Additionally, and possibly related, by providing a window of enhanced temporal coordination (most likely in the source population, i.e., the OB), the respiratory rhythm would favor spike transmission. This would occur because synchronous inputs have a higher probability of firing a target neuron than asynchronous inputs.^9,59,60^ Hence, we suggest that the combination of subspace communication and temporal coordination might be a canonical strategy to facilitate corticocortical interactions.

### Directionality in the canonical olfactory pathway

Long-range loops play a crucial role in various sensory and motor systems across the brain.^61,62^ In olfaction, the primary long-range pathway is formed by the connection from the OB to the PCx, although other significant loops exist (e.g., OB to anterior olfactory nucleus).^19,22,45,50^ Traditionally, these pathways are conceptualized in terms of their information flow^8,12,63^ as either feedforward (proceeding from the lower sensory area to a higher area, as in the case of OB→PCx) or feedback (going from the higher to the lower sensory area, as in the case of PCx→OB). Our results demonstrate that while feedforward interactions are predominant, feedback also occurs in the OB-PCx communication subspace, constituting a genuine loop. Notably, feedforward communication primarily occurred during inhalation, while feedback took place in a brief window during exhalation.

Furthermore, by analyzing how the local recurrent PCx circuit affects CS activity, we were able to show that PCx activity inhibits OB neurons (Fig. 5B, inferred from TeLC recordings showing a higher sustained OB response compared to control), consistent with previous reports showing that the excitatory centrifugal PCx output contacts granule cells within the OB and produces inhibition.^19,21^ From a functional perspective, previous results showed that PCx inhibition into OB decorrelates odor representations in this area and changes the tuning properties of single cells.^21^ This mechanism could potentially explain why OB-PCx spontaneous CS correlations decrease during odor stimulation. Under this scenario, during odor presentations PCx feedback changes OB activity, thereby promoting communication through the odor specific channel and decreasing communication through the spontaneous CS.

### Spontaneous vs. evoked activity and the representation of odor in the main olfactory pathway

How does spontaneously organized neural patterns relate to stimulus-driven activity? Through an examination of spontaneous communication and subsequent analysis of how these patterns change with sensory stimuli, we demonstrate that neuronal populations exhibiting spontaneous correlation also transmit odor information during sensory stimulation. Additionally, we observed that certain neuronal groups seem to communicate more robustly during odor stimulation, evidencing the recruitment of additional neurons during olfaction, a result evidenced more strongly in the PCx, where the odor CS contained more odor identity information than the spontaneous CS.

A possible interpretation of these results is that spontaneously correlated neurons form a fundamental scaffold for olfaction, providing a “good-enough” odor representation. On the other hand, neurons that communicate exclusively during odor stimulation contribute finer-grained details, enriching the overall representation. In support of these ideas, data from flies and mice shows that odor encoding populations fall between two extremes, either exhibiting reliable firing patterns across trials or displaying high trial-to-trial variability.^64^ The former property potentially adds robustness of the code while the latter might add discriminatory power.

### Methods

We analyzed simultaneous OB and PCx recordings generously made available by Bolding and Franks through the Collaborative Research in Computational Neuroscience data-sharing website (http://crcns.org, pcx-1 dataset). The SIMUL and TeLC-THY experiments were employed. Detailed descriptions of the experimental procedures are found in previous publications.^22,23,65^ The Institutional Animal Care and Use Committee of Duke University approved all protocols. Below we describe the analytical methods employed and the relevant experimental procedures from the original publication.

### Animals

Adult mice were employed (>P60, 20–24 g); animals were housed in single cages on a normal light-dark cycle. For the optogenetic experiments, the mice employed were adult Thy1ChR2/ChR2-YFP, line 18 (Thy1-COP4/EYFP, Jackson Laboratory, 007612). For the combined optogenetics and TeLC expression experiments, adult offspring of Emx1Cre/Cre mice crossed with Thy1ChR2/ChR2-YFP mice were employed.

### Adeno-associated viral vectors

For the TeLC experiments, AAV5-DIO-TeLC-GFP was expressed under CBA control (6/7 mice) or synapsin (1/7 mice); since effects were similar in either, data were pooled together. Three 500 nL injections in the PCx (AP, ML, DV: +1.8, 2.7, 3.85; +0.5, 3.5, 3.8; –1.5, 3.9, 4.2 mm; DV measured from brain surface) were employed to achieve TeLC expression. All recordings took place ∼14 days after the promoter injection. All viruses were obtained from the University of North Carolina-Chapel Hill (UNC Vector Core).

### Data acquisition

The electrophysiological signals were recorded through 32-site polytrode acute probes (A1x32-Poly3-5mm-25 s-177, Neuronexus) with an A32-OM32 adaptor (Neuronexus) through a Cereplex digital headstage (Blackrock Microsystems). Data were acquired at 30 kHz, unfiltered, employing a Cerebus multichannel data acquisition system (BlackRock Microsystems). Respiration was acquired at 2 kHz by analog inputs of the Cerebus system. The respiration signal was measured employing a microbridge mass airflow sensor (Honeywell AWM3300V), which was positioned opposite to the animal’s nose. Inhalation generated a negative airflow and thus negative changes in the voltage of the sensor output.

### Electrode and optic fiber placement

For OB recordings, a Patchstar Micromanipulator (Scientifica) holding the silicon probe was set to a 10-degree angle in the coronal plane to target the ventrolateral mitral cell layer. The probe was first positioned above the OB center (4.85 mm AP, 0.6 mm ML from bregma) and then lowered following this angle until dense spiking activity was found from the mitral cell layer, typically between 1.5 and 2.5 mm from the OB surface. For PCx recordings, the micromanipulator was employed to position the recording probe in the anterior PCx (1.32 mm AP and 3.8 mm ML). Recordings were targeted 3.5–4 mm ventral from the brain surface at this position and were further adjusted according to the LFP and spiking activity monitored online. The electrode sites spanned 275 µm along the dorsal-ventral axis. The probe was lowered until an intense spiking band was found, which covered 30–40% of electrode sites near the correct ventral coordinate, thus reflecting the piriform layer II. For the optogenetic experiments stimulating OB cells in Thy1ChR2/ChR2-YFP mice, the optic fiber was placed <500 µm above the OB dorsal surface.

### Spike sorting and waveform characteristics

Spyking-Circus software was employed to isolate individual units (https://github.com/spyking-circus). All clusters which had more than 1% of ISIs violating the refractory period (<2 ms) or appearing contaminated were manually removed. Units that showed similar waveforms and coordinated refractory periods were merged into single clusters.

### Odor delivery

Stimuli consisted of monomolecular odorants diluted in mineral oil. The odorants employed were: hexanal, ethyl butyrate, ethyl acetate, 2-hexanone, isoamyl acetate, and ethyl tiglate. Odor stimuli were presented for one second through an olfactometer controlled by MATLAB scripts and repeated every ten seconds.

### Anesthesia recordings

In 11 out of 13 mice, a combination of ketamine and xylazine at a dose of 100/10 mg/kg was injected intraperitoneally, leading to consistent anesthesia for 30–45 minutes. Body temperature was regulated using a heating pad throughout this period. The initial 30 minutes after the onset of slow oscillation were utilized for all assessments. One mouse was excluded from the analysis due to the absence of any physiological indications of anesthesia.

### Analytical Methods

For all analyses, we used Python 3 with numpy (https://numpy.org/), scipy (https://docs.scipy.org/), matplotlib (https://matplotlib.org/), sklearn (https://scikit-learn.org/stable/), and pyinform (https://elife-asu.github.io/PyInform/).

### Individual cell spiking preprocessing

Only putative principal OB or PCx cells were employed. The spike times were rounded to match the respiration signal resolution (0.5 ms or 2000 Hz sampling rate). Then, we convolved the spike times of each unit with a 10 ms standard deviation Gaussian kernel employing the convolve scipy function (in close similarity with the original publication), which gave rise to a smoothed spiking activity. To obtain the MUA signals, we summed the smoothed firing rate of each individual neuron within a population, obtaining a 1-d signal.

### Canonical correlation analysis to study the communication subspace

Before performing canonical correlation analysis (CCA), all smoothed spike trains were z-scored by subtracting the mean firing rate and dividing the result by the standard deviation. CCA seeks to find the sets of weights *Wx* and *Wy* that maximize the Pearson correlation coefficient between the *X* and *Y* vectors (when weighted by *Wx* and *Wy*). In our case, *X* and *Y* correspond to the smoothed firing rates of two different neuronal populations. CCA is thus defined by:

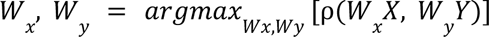

where:

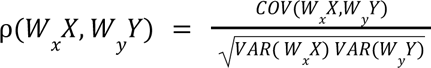

*COV*() and *VAR*() being the covariance and variance respectively.

*Wx* and *Wy* are the weight vectors that point-by-point multiply *X* and *Y* activity (i.e., *W_x_X* = *W_x1_x*1 + *W_x2_ x*2 + … + *W_xN_xN*). They are related to the *X* and *Y* firing vectors through their Σ*xy* covariance matrix:

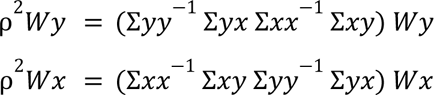

We can see that the above equation is in the form of the standard eigenvalue problem *Av* = λ*v*, where λ = ρ is the eigenvalue of the *A* = Σ*yy* Σ*yx* Σ*xx* Σ*xy* matrix, and *Wy* = *v* is the corresponding eigenvector. Thus, the *Wy* and *Wx* vectors are found as the eigenvectors of the Σ*yy*^−1^ Σ*yx* Σ*xx*^−1^ Σ*xy* and Σ*xx*^−1^ Σ*xy* Σ*yy*^−1^ Σ*yx* modified covariance matrices respectively.

Note that there exist *M Wy* and *Wx* vectors that satisfy the previous equations, which are all the eigenvectors of the above mentioned covariance matrices, where *M* = *min*(*Nx*, *Ny*), that is, the minimum number of neurons in either the *X* or *Y* population. Each *M* vector pair defines one communication subspace (CS) pair. These pairs are ordered according to the correlation coefficient (R) they produce, i.e., *R*(*CS*_1_) > *R*(*CS*_2_) > *R*(*CS*_3_) >… > *R*(*CS*_*M*_). Of note, because each CS pair is one of the eigenvectors of the modified covariance matrix, they are orthogonal against each other, implying that the different CS dimensions for a given area are not correlated.

For the practical implementation, we employed the *CCA* sklearn function employing *max_iter=100000*, *tol = 1e-12.* The weights were obtained through the *x_weights_* and *y_weights_* methods of the *CCA* object. The CS activity was obtained through the *fit.transform CCA* method. CS and MUA signals were z-scored to allow comparisons.

### Time-lagged CCA

The correlation between CS activities was computed using the Pearson correlation coefficient *pearsonr* stats.scipy function. To find the optimal lag between both OB and PCx CS signals, we temporally lagged PCx activity by different times and computed CCA to find the time-lag which maximizes the OB-PCx CS correlation. The weights obtained for this time-maximizing correlations were employed subsequently to obtain the respective CS signals.

### Surrogate analysis

To understand how many significant CS pairs exist, we contrasted CS correlations through a surrogate analysis. To generate a surrogate distribution to compare CS correlation values, we circularly shifted the spiking activity of one of the populations in time by random amounts. For each surrogate data, we computed the CCA and obtained the correlation coefficient for the first most correlated pair (CS_1_). The procedure was repeated 100 times for each session, and all surrogate correlation coefficients were pooled together to yield a N = 1300 surrogate distribution (100 surrogates from 13 sessions across 12 animals). We then assessed the significance by comparing the correlation coefficient for each CS pair against the null surrogate distribution. Only correlation values above the 99th percentile were deemed significant (p<0.01).

### Cross-correlation analysis

Cross-correlations between OB-PCx CS signals or MUA signals were estimated employing the *correlate* signal.scipy function using the *mode=’same’* and *method=’fft’* parameters.

### Directionality analysis

We calculated the transfer entropy between the binarized CS_1_ signals to study the directionality between OB and PCx CS_1_ signals. The binarization procedure assigned a 0 when the CS_1_ signal was below the mean CS_1_ or a 1 when the CS_1_ signal was above the mean. Then, transfer entropy was estimated using a 10 ms history length between signals.

Specifically, transfer entropy is defined as:

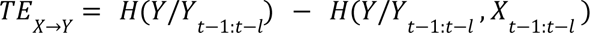

where *H*(*Y*/*Y*_*t*−1:*t*−*l*_) is the conditional Shannon entropy of the *Y* signal given its previous history. *l* = 10 ms in our analysis (20 time points).

The same analysis was employed to study Resp-CS_1_ directionality, the only difference being that Resp and CS_1_ signals were down-sampled to 40 Hz and *l* = 500 ms in our analysis (20 time points).

### Principal component analysis

PCA finds subsets of correlated neurons within each area. The principal components (PCs) are found through the eigenvectors (*v*) of the covariance matrix (Σ*xx*), Σ*xx v* = λ *v*, where λ are the eigenvalues (explained variance of that PC) of Σ*xx*. We employed the *PCA* sklearn function to compute the PCs of the OB and PCx activity (the same OB and PCx activity vectors employed for CCA were used for PCA).

### Power spectrum and coherence

We computed Welch’s modified periodogram to estimate the CS spectra (*welch* scipy function). Specifically, we employed a 1 s moving window with half a window overlap, setting the numerical frequency resolution to 0.1 Hz (by setting the nfft parameter to 10 times the sampling rate). Similarly, we computed the magnitude squared coherence (OB-Resp, PCx-Resp, OB-PCx) through the *coherence* scipy function employing the abovementioned parameters.

To obtain a surrogate CS-Resp coherence distribution (Fig. S4B), we circularly shifted the Resp signal in time by random amounts and repeated the procedure 100 times per session. Individual CS-Resp coherence values were then compared against the null distribution (made from CS_1_-shuffled Resp coherence), and only the values above the 99th percentile were considered significant (p<0.01).

### Spike-phase, correlation-phase, and directionality-phase plots

We computed the average spiking activity per respiration phase by first band-pass filtering the respiration signal using the *eegfilt* function (Delorme and Makeig, 2004) adapted for Python 3 (available at https://github.com/joaqgonzar/Gamma_Oscillations_PCx). We filtered the signal between 1–3 Hz to obtain the slower frequency component. The phase (angle) of the filtered respiration signal was estimated from their analytical representation based on the Hilbert transform. We then binned the phase time series into 300 bins and computed the mean spiking activity of each neuron for each respiration phase bin. Similarly, for each respiration phase bin, the correlation between the OB CS_1_ and PCx CS_1_ was computed using only the time points associated to it. For the directionality-phase plot, we computed transfer entropy for each phase bin employing a history length of 10 ms (see transfer entropy details above).

### Spike-spike histogram

For each recording session, we measured spike-spike histograms between OB and PCx neurons. We first selected the ten most positive OB neurons according to their CS_1_ weights and the ten most positive and negative CS_1_-weighted PCx neurons. We constructed histograms (employing 10 ms bins) of the time differences between an OB neuron spike and the spikes of a PCx neuron. To compare the shape of the histograms (Fig. 4D), we z-scored them by subtracting the mean and dividing them by the standard deviation. Peak activity (excitation/inhibition) shown in Fig. 4E was obtained by selecting from each normalized histogram the time lag that had the largest positive (excitation) or negative (inhibition) value.

### Odor decoding from CS activity

We employed a supervised linear classifier to decode either odor identity (six different odorants at 0.3% v./v. concentration) or odor concentration (four concentrations from 2 different odorants) from the CS activity. We supplied the average CS activity (CS_1_, [CS_1_,CS_2_], [CS_1_,CS_2_,CS_3_] etc.) during the first 500 ms following the inhalation start. The classification algorithm was a linear support vector machine with a stochastic gradient descent optimization, implemented using the *sgdclassifier* sklearn function. Each session was trained and tested separately using leave-one-out cross-validation through the *LeaveOneOut* sklearn function; for each iteration, a single odor trial was withheld from training and tested later; the procedure was repeated for all odor trials, and the mean accuracy was reported.

### Toy model for studying cross-area communication

We employed two different strategies to construct a toy model to validate CCA:

1. We generated two 10×10000 random firing rate matrices (X and Y areas in Fig. S1), representing ten neurons in each area and 10000 time points. On top of that, we imposed several activation times (specific moments in time when firing rates increased substantially). These activation times could be either local (occurring in only one matrix) or shared between both matrices (Fig. S1A,B). To study lagged communication (Fig. S3), we made the common activation times (shared between both matrices) to be lagged in time by two time points. To overcome the fact that CCA cannot deal with lagged correlations, we either convolved the matrices with a Gaussian kernel or lagged one of the two matrices in time before performing CCA.
2. We also tested a synthetic case in which a set of Y neurons would receive a linear combination of X outputs (without any common activation times). In the example of Fig. S1C,

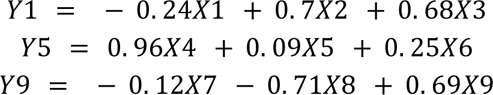 Where Y1 stands for neuron 1 of the Y area and so on. The weights of the combinations were picked to be orthogonal vectors to make the different combinations independent. For Figure S1D, we increased the number of independent combinations from 0 to 10 and measured the correlations obtained from each CS activity pair.

### Statistics

We show group data as either mean ± SEM or regular boxplots showing the median, 1st, 3rd quartiles, and the distribution range without outliers. We employed paired and unpaired t-test to compare between groups. For Figures 5D and 7E we employed two-way ANOVAs. We set p<0.05 to be considered significant (in the figure panels, *, ** and *** denote p<0.05, p<0.01, and p<0.001, respectively).

## Data availability

All the data employed is freely available at http://dx.doi.org/10.6080/K00C4SZB. Previously Published Datasets: Bolding and Franks, 2018b Collaborative Research in Computational Neuroscience (http://crcns.org/). ‘Simultaneous extracellular recordings from mice OB and PCx and respiration data in response to odor stimuli and optogenetic stimulation of OB‘ (https://crcns.org/data-sets/pcx/pcx-1). The original codes employed are available at: https://github.com/joaqgonzar/CS_OB-PCx.

## Acknowledgements

We thank Diego Laplagne and Vitor Lopes-dos-Santos for their critical reading of our manuscript. JG was supported by Comision Academica de Posgrado (CAP), Programa de Desarrollo de Ciencias Básicas (PEDECIBA), and Comisión Sectorial de Investigación Científica (CSIC). PT was supported by PEDECIBA and CSIC. KAB was supported by the Monell Chemical Senses Center. ABLT was supported by Conselho Nacional de Desenvolvimento Científico e Tecnológico (CNPq), Coordenação de Aperfeiçoamento de Pessoal de Nível Superior (CAPES), and the Alexander von Humboldt Foundation.

## Author Contributions

JG: Conceptualization, Software, Formal analysis, Visualization, Methodology, Writing – original draft, Writing – review and editing. PT: Supervision, Funding acquisition, Writing – review and editing. KAB: Investigation, Resources, Data Curation, Writing – review and editing. ABLT: Supervision, Funding acquisition, Project administration, Writing – review and editing.

## Supplementary Material

**Figure S1.**
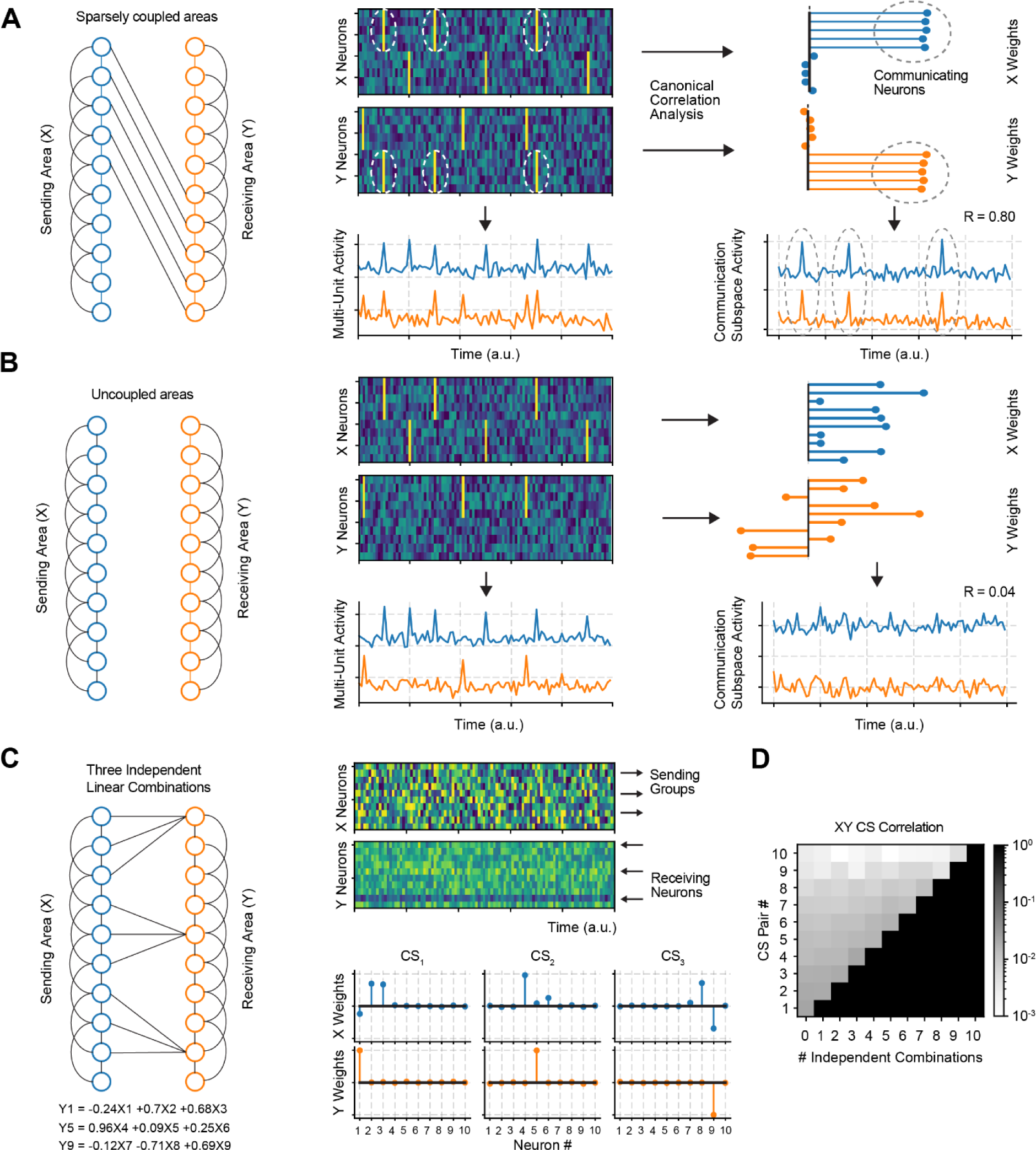
Canonical correlation analysis unveils embedded patterns of communication between two neuronal populations. **A** Toy example of 2 areas (X,Y) communicated instantly through common activation times in neurons 1-5 (X area) and 5-10 (Y area). Activation times are imposed on top of random, independent activity in each area. In addition to the common activation times, we also included independent activation times (not shared between both areas). To unveil the correlation present in this example, we employed CCA, which correctly revealed the neurons in both areas which were activated simultaneously. **B** Same case as **A,** except the spikes from neurons 1-5 (X area) are not communicated to area Y. **C** Toy example of 2 areas (X, Y) communicated instantly by three independent linear combinations of neurons (as shown by the equations below the leftmost panel). In this case, no activation times are present, and each neuron exhibits random firing rate fluctuations in each area. On top of that, neurons 1,5,10 of area Y receive a linear combination of the activity of three X neurons. CCA correctly identifies the linear combinations present in the data. **D** Number of correlated canonical pairs plotted as a function of the number of independent linear combinations present in the X and Y data. Notice that when only one independent combination is present, only one canonical pair shows significant correlation.

**Figure S2.**
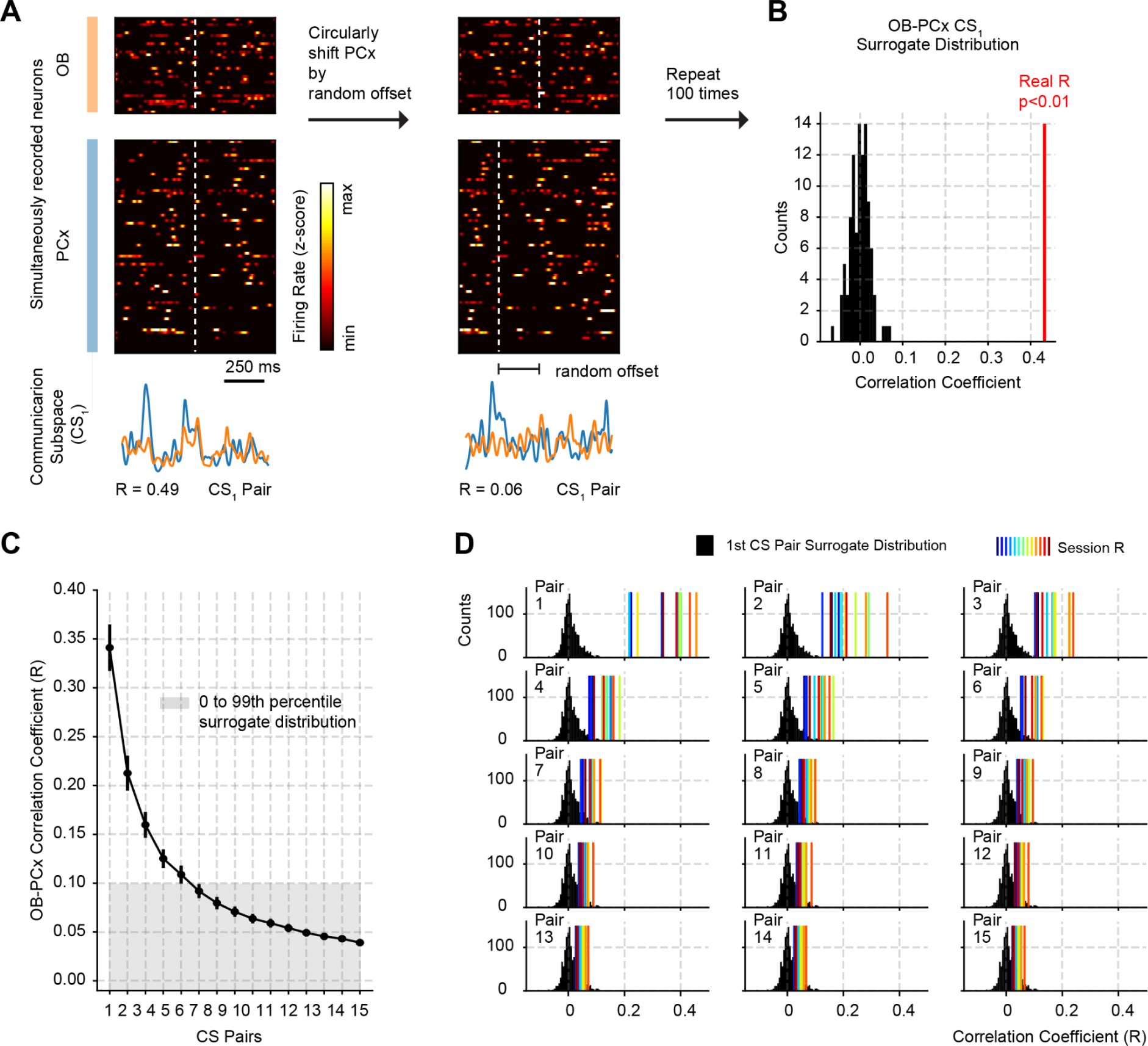
Characterizing correlations across communication subspace pairs. **A** Example showing the surrogate construction method. Raster plots show the normalized (z-scored) firing rate of each neuron in OB and PCx. The plot at the bottom shows the CS_1_ activity for each area (1st pair) and its corresponding correlation coefficient (R) value. To obtain a surrogate, a circular time shift is carried on the PCx population before computing the CS_1_; the R is then recomputed. **B** The procedure is repeated 100 times to yield a surrogate CS_1_ OB-PCx R distribution. The real R value is shown in red. **C** Average Pearson correlation coefficient (R) for each CS pair (± SEM, n = 13 recording sessions from 12 mice). The gray area shows the 0 to 95th percentile range of the surrogate R distribution. For each session, 100 surrogates were computed by randomly circularly shifting one of the population vectors (in this case, the PCx), and then computing the canonical correlation and obtaining the R of the first pair. All 100 coefficients for each animal and session (n = 13 recording sessions from 12 mice, 1300 surrogate values total) were pooled together, and the corresponding percentiles were computed. The procedure was modified following the method employed by ^13^. **D** R values for each session (colored lines) plotted along the surrogate R distribution. Each plot shows the R for one communication subspace pair. Note that the surrogate distribution is the same across panels, as the surrogate R of the first CS was employed for all pairs.

**Figure S3.**
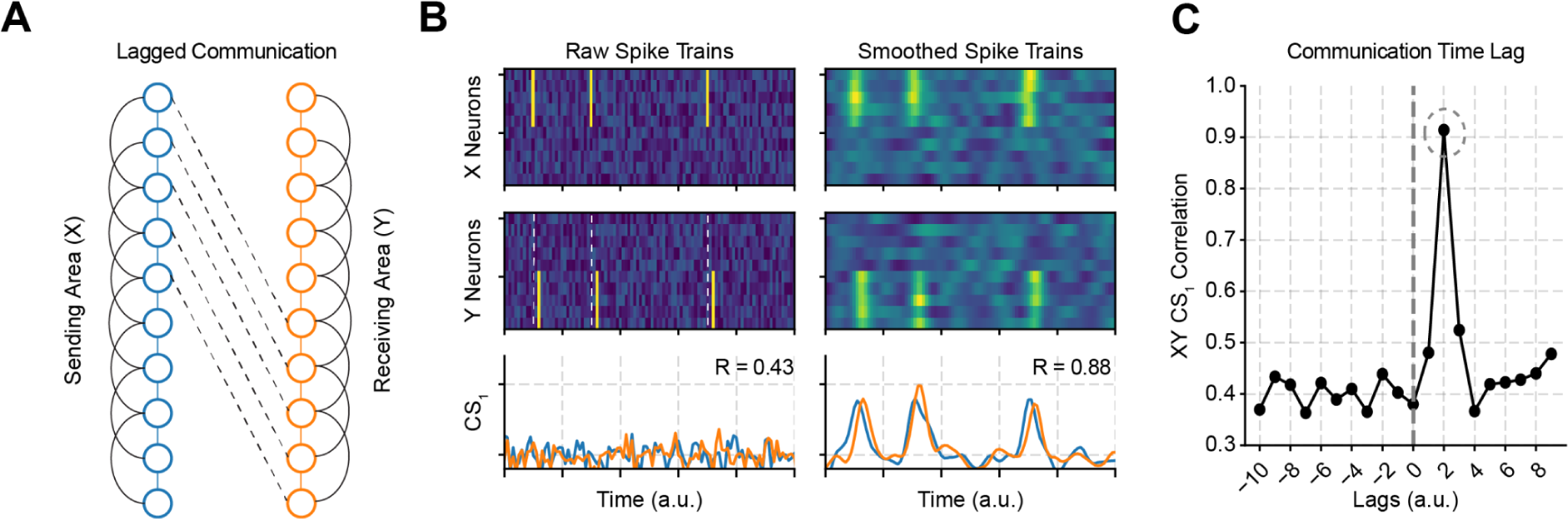
Capturing lagged communication through canonical correlation analysis. **(A,B)** Toy example of 2 areas (X,Y) communicated through lagged activation times between neurons 1-5 (X area) and 5-10 (Y area). Note in **B** left that the activation times of X and Y populations do not coincide and that a convolution with a Gaussian kernel smooths the activity and unveils a common pattern observed in the projected CS_1_ activity. **C** As an alternative, the lagged communication can also be revealed with no smoothing by lagging the activity of one population against the other prior to computing the CS weights. The dashed circle highlights the CS_1_ pair with highest correlation.

**Figure S4.**
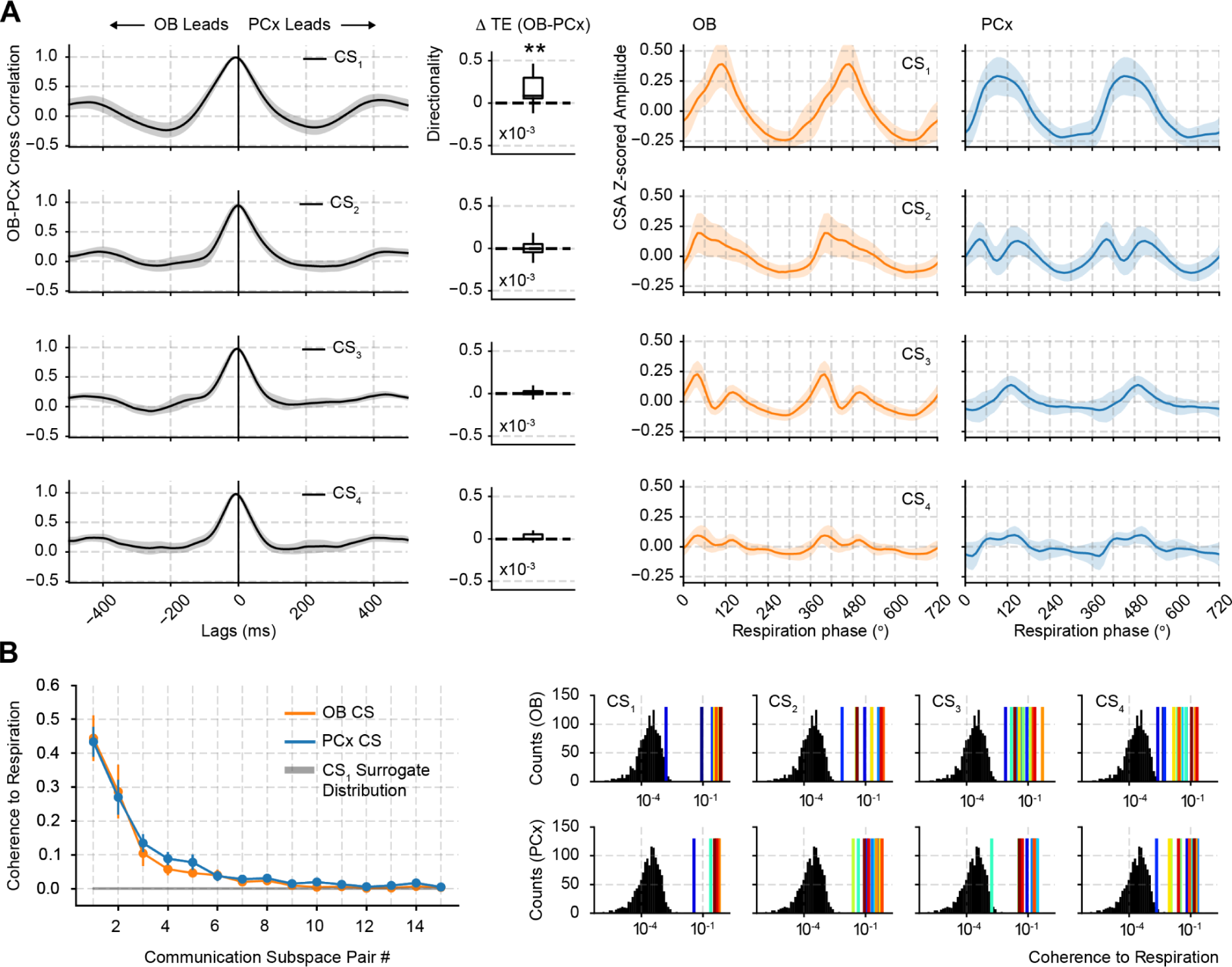
Directionality and respiratory entrainment for the top four unlagged CS pairs. **A** Left: Average OB-PCx communication subspace cross-correlogram for the top four unlagged CS pairs (i.e., the CS weights were obtained without lagging the spike time series by 25 ms) and directionality analysis (Transfer Entropy) between both areas. The difference (delta) between OB→PCx and PCx←OB transfer entropy estimates is plotted for each signal. Boxplots show the median, 1st, 3rd quartiles, the minimum, and the maximum (excluding outliers); n = 13 recording sessions from 12 mice. ** p < 0.01. Right: Average CS of each pair as a function of respiratory phase (± SEM; n = 13 recording sessions from 12 mice). **B** Left: CS-Respiration coherence (at 2.5 Hz, the breathing rate) as a function of the CS pair number. The gray line shows the 95 percentile of the surrogate distribution. Surrogates were computed by circularly time shifting the respiratory activity and computing the CS_1_-Resp coherence. Right: CS-Resp Coherence values for each session (colored lines) plotted along the surrogate Coherence distribution. Each plot shows the coherence for one communication subspace pair.

**Figure S5.**
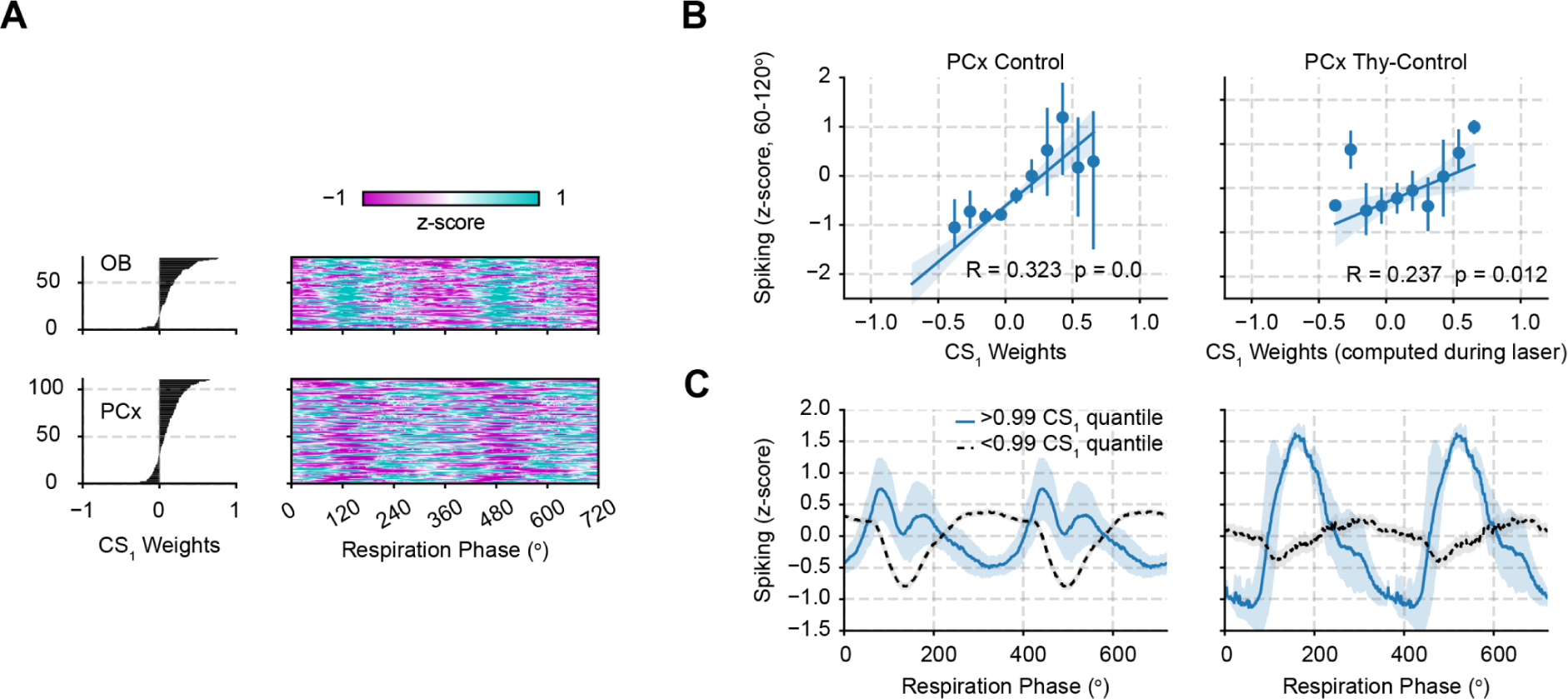
Laser-evoked CS weights predict single cell respiratory spiking phase preferences. **A** Putative principal cell spiking during each phase bin of the respiration cycle (bottom; 0 degree corresponds to the start of the inhalation); neurons are sorted according to the canonical correlation weights (shown on the left panel). **B** Correlation between CS weights and CS_1_ activity during the 60-120 respiration phase. Error bars show the mean ± SEM binned CS. **C** Average spiking activity from single PCx units during the respiratory phase. Units were sorted by their CS_1_ weights either above or below the 99th quantile. Notice the anti-phase pattern exhibited by neurons in both categories.

**Figure S6.**
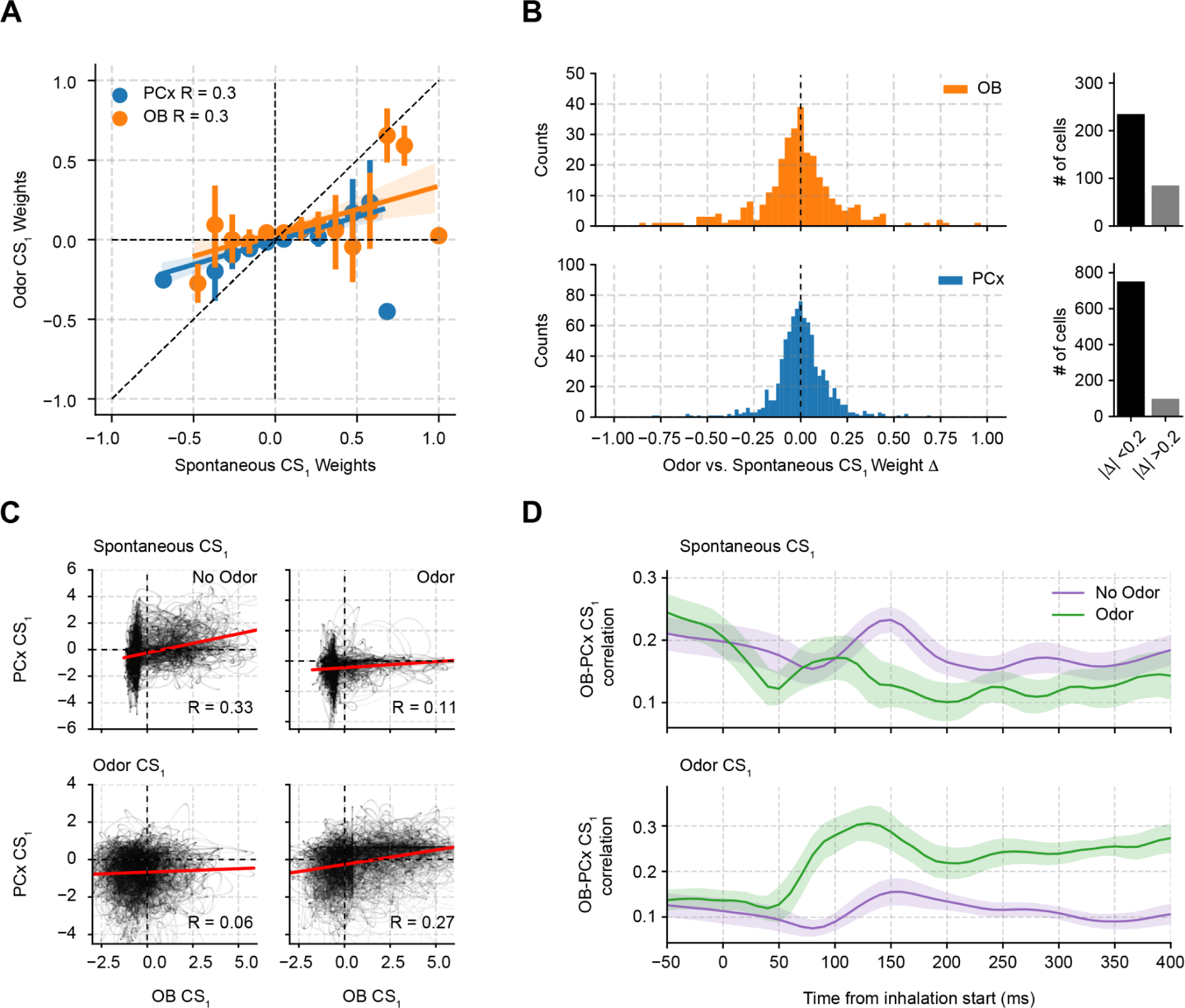
Spontaneous vs. odor-evoked CS activity and correlations. **A** Correlation between CS_1_ weights exclusively computed during odor and odorless periods. **B** Left: CS_1_ weight difference (delta) between CS computed during odor and odorless periods. Right: Bar plot showing the number of neurons in each area exhibiting CS_1_ weight absolute differences above and below 0.2. **C** OB-PCx CS_1_ correlation from a representative animal during odor and odorless cycles. For the top panel, CS_1_ weights were computed during spontaneous sniffs (no odors presented), while for the bottom panel CS_1_ weights were computed during odor sniffs. **D** Average OB-PCx CS_1_ correlation time-courses as a function of time from inhalation start (± SEM; n = 13 recording sessions from 12 mice). To compute correlations, we pooled all OB-PCx CS_1_ activity into 10 ms bins starting 50 ms prior to inhalation start. For each bin, the correlation coefficient for each session and animal was computed.

**Figure S7.**
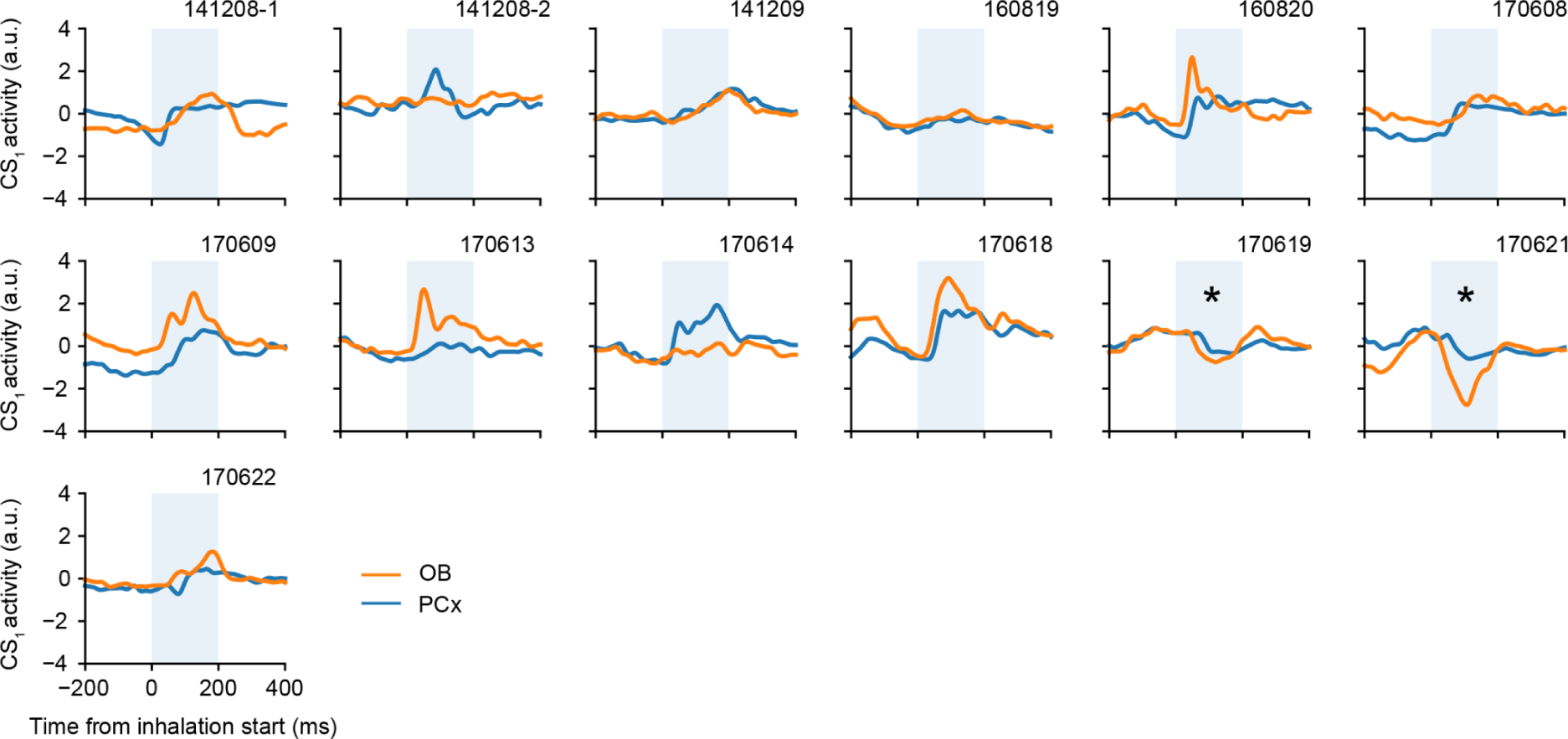
CS response to odors for each animal. CS_1_ activity (computed only during the presentation of odors) is plotted as a function of time from the inhalation start. Note that 0 time corresponds to the beginning of an inhalation where an odorant is present. The asterisks show two animals which exhibited a CS_1_ activity decrease in response to odors. To normalize these traces, we inferred whether the response was positive or negative based on the sign of the maximum response (during the 200 ms window showed by the shaded area) compared to the sign of CS_1_ activity at time zero.

**Figure S8.**
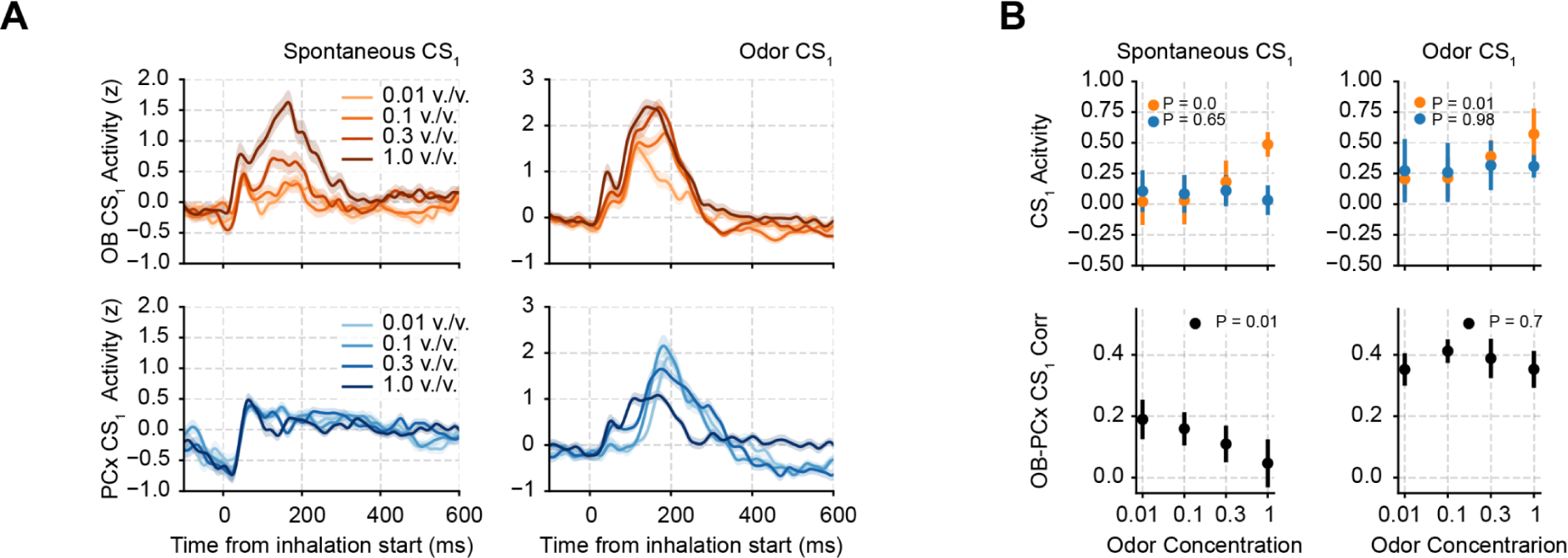
Odor CS_1_ correlations show concentration invariant responses to odorants. **A** Average CS_1_ activity triggered by inhalation start during the presentation of a single odorant at increasing concentrations (± SEM; n = 120 trials from 5 mice). CS_1_ weights were computed either employing only odorless periods (spontaneous CS) or only during the presentation of odors (Odor CS). **B** Top: Average CS_1_ amplitude as a function of odorant concentration (± SEM; n = 5 mice). Bottom: Average OB-PCx CS_1_ correlation as a function of odorant concentration (± SEM; n = 5 mice).

**Figure S9.**
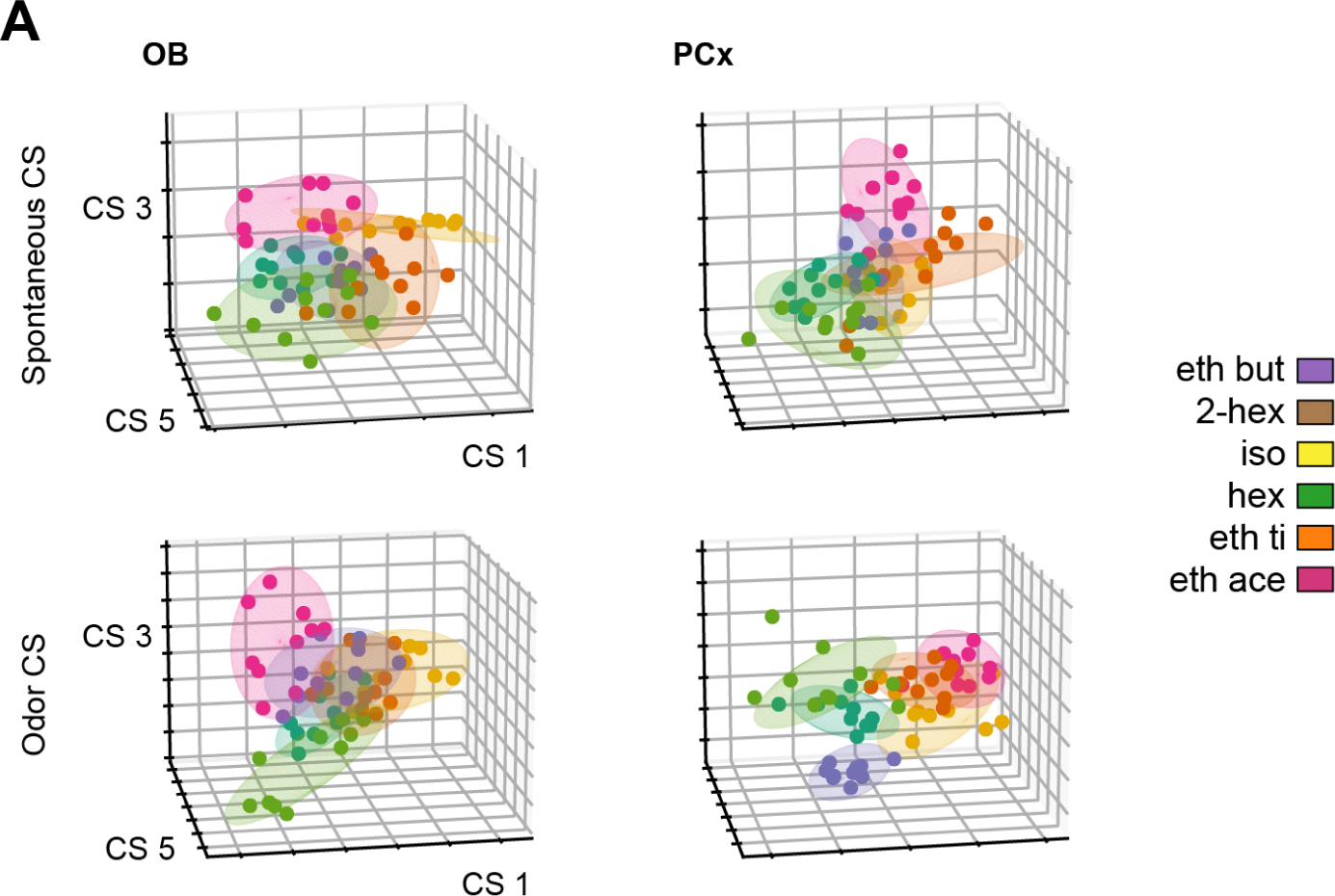
CS odor clusters for six monomolecular odorants. Example CS activity for 3 CS pairs from a representative animal obtained during the presentation of 6 different odorants. Each dot shows the average CS activity (0-500 ms following inhalation start) during an individual trial. Ellipses show the 95% confidence interval for each odor.

**Figure S10.**
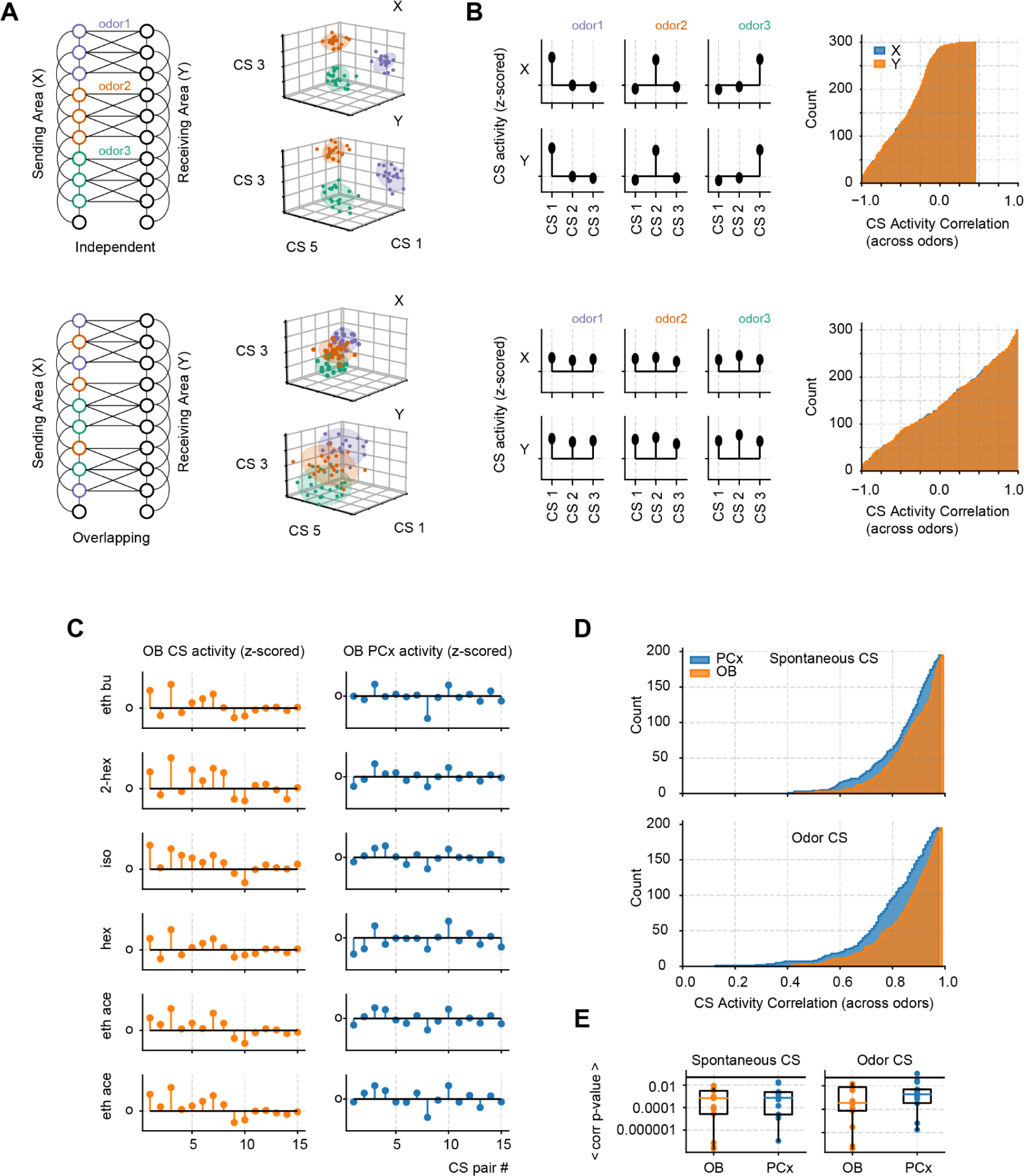
CS_1_ odor transmission involves overlapping neuronal assemblies. A Left: Toy-model example of 2 areas (X and Y) transmitting odor information through non-overlapping (top) and overlapping (bottom) combinations of neurons. Each odor produced the instant activation of the neurons representing it on the X population. Note that the connectivity between the X and Y areas remains identical in both examples, only the neurons responding to odors in X are changed. Right: CS activity for 3 CS pairs for all odor activations. **B** Left: Average CS activation profiles for each odor. We plot the mean activity (across all trials) for each CS component in each area, thus obtaining a measure of how each CS is activated by each odor. Right: Distribution of correlation coefficients comparing the CS activation profiles. We correlated the CS profiles across odorants. For example, in the case of three odors: CS[odor 1] vs. CS[odor 2]; CS[odor 1] vs. CS[odor 3]; CS[odor 2] vs. CS[odor 3]. Then the distribution of all correlation values is plotted. Notice much higher correlation when odor coding takes place by overlapping combinations of neurons (bottom panel). **C** Average CS activation profiles for the actual OB-PCx data employing six different odorants. The top 15 Odor CS pairs of a representative animal are plotted. **D** Distribution of correlation coefficients comparing the CS activation profiles in the OB-PCx data. All sessions and animals were pooled together. Both the spontaneous and Odor CS distributions are shown. Note high correlation coefficient values in the real data, similar to the simulated data shown in the bottom panel of B. **E** Boxplots showing the average p-values (n = 13 recording sessions from 12 mice) associated with the correlation coefficients for the CS odor activation correlations.

## Supplementary results

**(Figure S10)**

Our toy-model results had shown that independent linear combinations could be captured by the activity of different CS pairs. Therefore, by studying the activation of the different CS pairs we could infer if the transmission of odorant identity involves independent or overlapping combinations of neurons. In the former case, modeling results show that each CS pair would capture an independent combination involving a single odorant, giving rise to isolated odor clusters in the CS space (i.e., the space defined by employing each CS pair as an axis). In the latter case, since neurons transmitting different odors overlap, each CS pair would not transmit a single odor, giving rise to overlapping clusters (Fig. S10A).

To further quantify the CS transmission overlap, we measured how each CS pair activates to each odor (“CS activation profiles”) by computing the average CS amplitude (for each CS pair) for each odor presentation. We quantified the similarity in the CS activation profiles by calculating the Pearson correlation of the CS activation profiles (i.e., CS_activity_odor1 = [1,2,3] vs. CS_activity_odor2 = [2,1,2]: R = 0.0). Modeling results show that in the independent transmission case, CS activation profiles for different odors do not exceed 0.5 R values (Fig. S10B, top row). On the other hand, for the overlapping case, CS activation profiles exhibit higher correlation values (Fig. S10B, bottom row).

Our OB-PCx results show that odor clusters do overlap in CS space (Fig. 7D and S10), thus resembling the overlapping combinations of neurons case in our toy model (Fig. S10A,B bottom panels). Moreover, we found that each odor activates several CS pairs in both areas (Fig. S10C). The activation of CS pairs was correlated across odors, exhibiting a large proportion of significantly correlated CS activation profiles (Fig. S10D,E; the distribution of all correlation values across sessions is plotted, note that each session contributes 15 correlation values, six different odorants). Thus, our modeling and experimental results suggest that the transmission of odorant identity occurs through mixed combinations of neurons, likely caused by overlapping odor representations in the early olfactory system.

